# *In vitro* pectate lyase activity and carbon uptake assays and whole genome sequencing of *Bacillus amyloliquefaciens* subsp. *plantarum* strains for a pectin defective pathway

**DOI:** 10.1101/2021.01.03.425148

**Authors:** Mohammad K. Hassan

## Abstract

The pectin lyase activity of 59 *Bacillus amyloliquefaciens subsp. plantarum (Bap)* strains was tested *in vitro* on Pectate Agar (PA) and Tris-Spizizen Salts (TSS) medium. Bap strains were cultured on TSA medium and washed three times with sterile water before the inoculation on PA media. Higher and lower pectate lyase activity were observed in six (AP193, AP203, AP299, AP80, AP102, and AP52) and four (AP 194, AP214, AP215, and AP305) *Bap* strains compared to other Bap strains. A total of 12 Bap strains (AP67, AP71, AP77, AP78, AP85, AP102, AP108, AP135, AP143, AP189, AP192, and AP193) grew vigorously on TSS medium. A total of six *Bap* strains (AP194, AP204, AP214, AP216, AP219, and HD73) had lower growth compared to other *Bap* strains. Pectin (1%) were used for *in vitro* PA and TSS medium. Pectate lyase and utilization activity were not found in *Bacillus thuringiensis* subsp. *kurstaki* strain HD73 compared to *Bap* strains. A draft genome sequence for strains AP194 and AP214 that were negative for pectin utilization were generated using an Illumina MiSeq. RAST analysis revealed that the pectin-associated gene altronate hydrolase *(uxaA)* absent in AP 214 strain. Multiple amino acid alignments of *exuT* and *uxuB* gene sequence showed dissimilarities among AP194, AP214, and reference *Bap* strains.

## 1. Introduction

Pectate lyase enzyme was first discovered in *Erwinia carotovora* and *Bacillus polymyxa* in 1962 (Starr, 1962) and has since been reported in many plant pathogenic and non-pathogenic bacteria such as *E. aroideae* (Kamimiya et al., 1977); *E. chrysanthemi* (Starr, 1972); *Clostridium multifermentas* (Macmillan, 1964); *Fusarium solani* (Crawford, 1987)*, B. amyloliquefaciens* (Fan X., 2008); and *B. subtilis* (Soriano et al., 2006). Mechanism of this enzyme activity in pathogenic and non-pathogenic bacteria could have a subtle difference in the mode of action. Pectate lyase breaks down polygalacturonate into D-galacturonate and D-glucuronate, making it available form for bacteria to use as a carbon source. Previous studies have reported that *B. subtilis* (Mekjian et al., 1999), *E. coli* K-12 (Nemoz et al., 1976), *E. carotovora* (Abbott, 2008), and *E. chrysanthemi* (Abbott, 2008) are capable of utilizing pectin as a sole carbon source and energy.

Whole genome sequencing is a useful method to find out the high resolution, base by base view, gene expression, and regulation of the strains (Anonymous, 2016). Rapid Annotation using Subsystem Technology (RAST) is designed to annotate the gene of complete prokaryotic genomes, and it uses the highest confidence first assignment that guarantees a high degree of genome consistency (Anonymous, undated). This study was designed to screen 59 Bap strains for pectate lyase and utilization of pectin as a sole carbon source. In addition, this study also included whole genome sequences analysis of two Bap strains AP194 and AP214 in figuring out their pectin defective genes. The purpose of this research was i) to screen a large collection of *Bap* strains for pectin degradation and utilization as a sole carbon source, ii) to conduct a comparative genomic analysis that includes *Bap* strains that lack the capacity for pectin utilization.

## 2. Material and methods

### 2.1. Bap strains preparation

A total of 59 Bap strains were streaked onto Tryptic Soy Agar (TSA) plates from the cryo stocks stored in the −80°C and incubated at 28°C for 24 hours. One loopful of bacteria was inoculated into 10ml TSB in a glass test tube and placed into a shaking incubator at 28°C overnight using 220 Revolutions Per Minute (RPM) for 24-48 hours. Three replicates were used for each of the Bap strains.

### 2.2. Pectate lyase activity test

The bacteria grew from cryo stocks in the −80°C on Tryptic Soy Broth (TSB) at 28°C overnight using 220 rpm for 5 ml culture. A one-ml aliquot was pipetted into the 1.5 ml microcentrifuge tube, and centrifugation was done for 5 minutes at 10,000 x g speed. The supernatant was discarded, and the process was repeated three times using sterile water. In the final bacterial pellet, 1 ml of the sterile water was added to a microcentrifuge tube and vortexed thoroughly to uniform the bacterial suspension. Then, it was transferred to 1 ml of the sterile water containing test tube to measure the turbidity of a bacterial suspension of all strains until the optical density at 600 nm was approximately 0.5. Twenty μl of this standardized bacterial suspension was used in triplicate onto pectate-agar (Pa) media (Kobayashi et al., 1999) to determine the pectin lyase activity. The pH 8.0 of 0.1M Tris-HCl buffer was adjusted for the medium and sterilized using 0.45 μm Nalgene syringe filter (Thermo Scientific, USA) separately. The pectate-agar media plates were incubated at 28°C for 24-48 hours and then 1% Cetyltrimethyl ammonium bromide (CTAB) was poured over the surface of the plate at room temperature.

### 2.3. Pectin carbon uptake test

The capacity of the 59 *Bap* strains to utilize as a sole carbon source were assessed using a minimal Tris-Spizizen salt (TSS) (Shingaki et al., 2003) as the base media supplemented with 1% plant pectin (citrus source) (Tokyo Chemical Industry Co., Ltd). The TSS media was filter-sterilized using a 0.2 μm polyethersulfone (PES) vacuum filter unit (VWR, USA) and adjusted the medium p^H^ 7.0 using 10N NaOH. Each of the bacterial cultures was prepared for the pectin lyase assays with bacterial suspensions washed three times in sterile water, normalized to an OD_600_ = 0.5, and then 100 microliters (μl) of a 1:100 dilution were used to inoculate 1.9 ml TSS + 1% pectin cultures to adjust the OD_600_ = 0.030, in triplicate. Broth cultures were incubated at 28°C with 220 rpm, and OD_600_ readings were reocorded over a 40 hr period. *Bap* strains AP 193 was used as the positive control and *Bacillus thuringiensis* subsp. *kirstaki* HD 73 was used as the negative control. *Bacillus thuringiensis* subsp. *kirstaki* HD 73 was obtained from the USDA-ARS culture collection (Ames, Iowa) that was identified as a non-pectin utilizing strain based on its genome sequence. This strain was not observed to growth using pectin as a sole carbon source.

### 2.4. Whole genome sequencing of Bap strains

The genomic DNA of AP194 and AP214 was extracted by E.Z.N.A. ^®^ DNA Isolation Kit and the DNA concentration was measured by Qubit^®^ 2.0 Fluorometer (Thermo Fisher Scientific, USA). Nextera DNA Library Preparation Kit (Illumina, Inc.) was used for Illumina MiSeq^®^ sequencing. A total of 50 ul reaction (3.75 μl genomic DNA, 25 μl TD buffer, 5 μl enzyme, and 16.25 μl nuclease free water) was prepared using Nextera sample prep kit. The prepared reaction was vortexed and centrifuged at 10,000x rpm before the incubation for 5 minutes at 55°C. Nextera PCR program cycle was used for PCR amplification using Eppendorf Thermal Mastercycler (Eppendorf AG, Hamburg). The holding temperature was 10°C. Zymo Research DNA Clean & Concentrator™-5 was used for purifying the tagmented DNA. A total of 250 μl DNA binding buffer was used for the 50 ul sample (5:1) and centrifuged for 30 seconds using 10,000 x g. A total of 200 μl DNA wash buffer was added and centrifuged it for 30 seconds. This process was repeated twice. The empty column was used for drying the sample and centrifuged for 2 minutes. A total of 25 μl DNA wash buffer was added and kept room temperature for one minute. Then, 50 μl reaction (20 μl tagmented DNA, 5 μl index 1, 5 μl index 2, 15 μl PCR master mix, and 5 μl PCR primer coctail) was prepared using Nextera prep kit. The temperatures for the PCR cycles were 12°C (3 min), 98°C (30 sec) for one cycle, 98°C (10 sec), 63°C (30 sec), and 72°C (3 min) for five cycles. The size-select purification kit was used for the cleaning of the 50 μl PCR products. The last step was a measurement of the DNA library concentration. DNA concentration was followed 16ng/μl, and 3.75μl genomic DNA were used for 60 ng concentration. Before running the MiSeq system, all instruments were cleaned using 1X Tween buffer. Then, all the samples were loaded to run the MiSeq^®^ system. After successful running the system, fastq.gz files were generated and saved as output files. Then, fastq.gz output files were imported, trimmed, and de novo assembled.

### 2.5. RAST analysis

Rapid Annotation using Subsystem Technology (RAST) version 2.0 was used for annotation of the whole genome sequences for strains AP194 and AP214. Fasta files of AP194 and AP214 were uploaded in RAST server. The annotated genome was viewed in a seed viewer link for AP194 and AP214.

### 2.6. CLC genomics analysis of *exuT* and *uxuB* protein gene sequence

Two pectin-associated genes *(exuT* and *uxuB)* sequences of 15 Bap reference strains were collected from the NCBI database. Each gene sequence was translated into an amino acid sequence using CLC Genomics Workbench 4.9 (CLC bio, Cambridge, MA, USA) software. Reference strains gene sequences were aligned with *exuT* and *uxuB* gene sequences of AP193, AP194, and AP214 strains.

## 3. Results

### 3.1. Bap strains preparation

A total of 59 Bap strains grew profusely within 24 hours in TSB and TSA plates. Most strains exhibited similar growth rate, but strains AP194, AP214, and AP52 grew slower than the rest. None of the Bap strains were cross contaminated.

### 3.2. Pectate lyase activity test

Clear zone appeared around bacterial colonies after 30 minutes. The magnitude of the zone of clearing was measured in milliliters (mm) and recorded in an Excel spreadsheet, with average zones of clearing determined for each of the *Bap* strains. The clear zone images of each plate were photographed using the AlphaImager^®^ HP high-performance imaging System. The pectinase clear zone diameter of each strain is shown in Table 1.

**Table 1.**
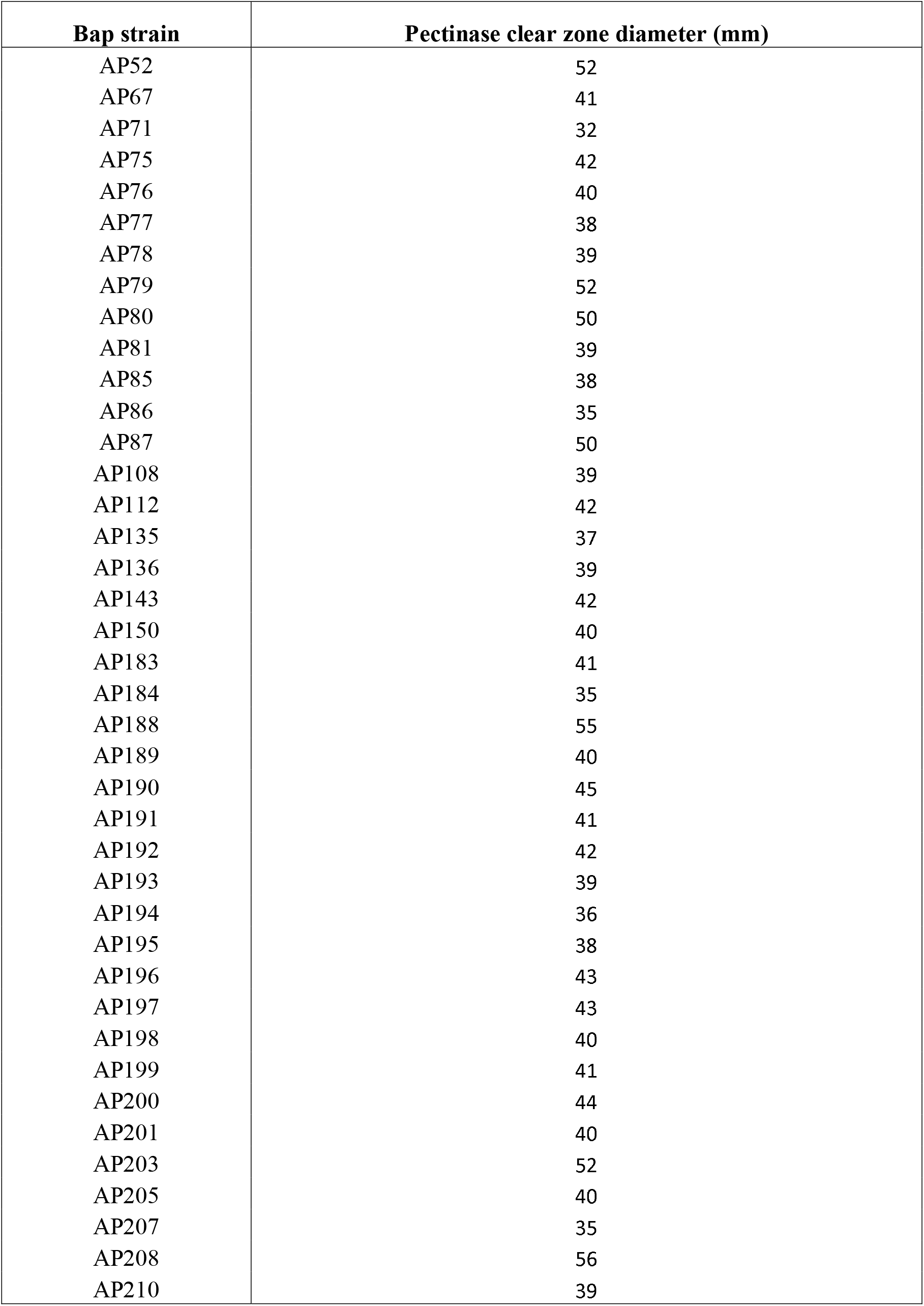
The clear zones range from 22 mm to 55 mm in Bap strains.

**Table 2.**
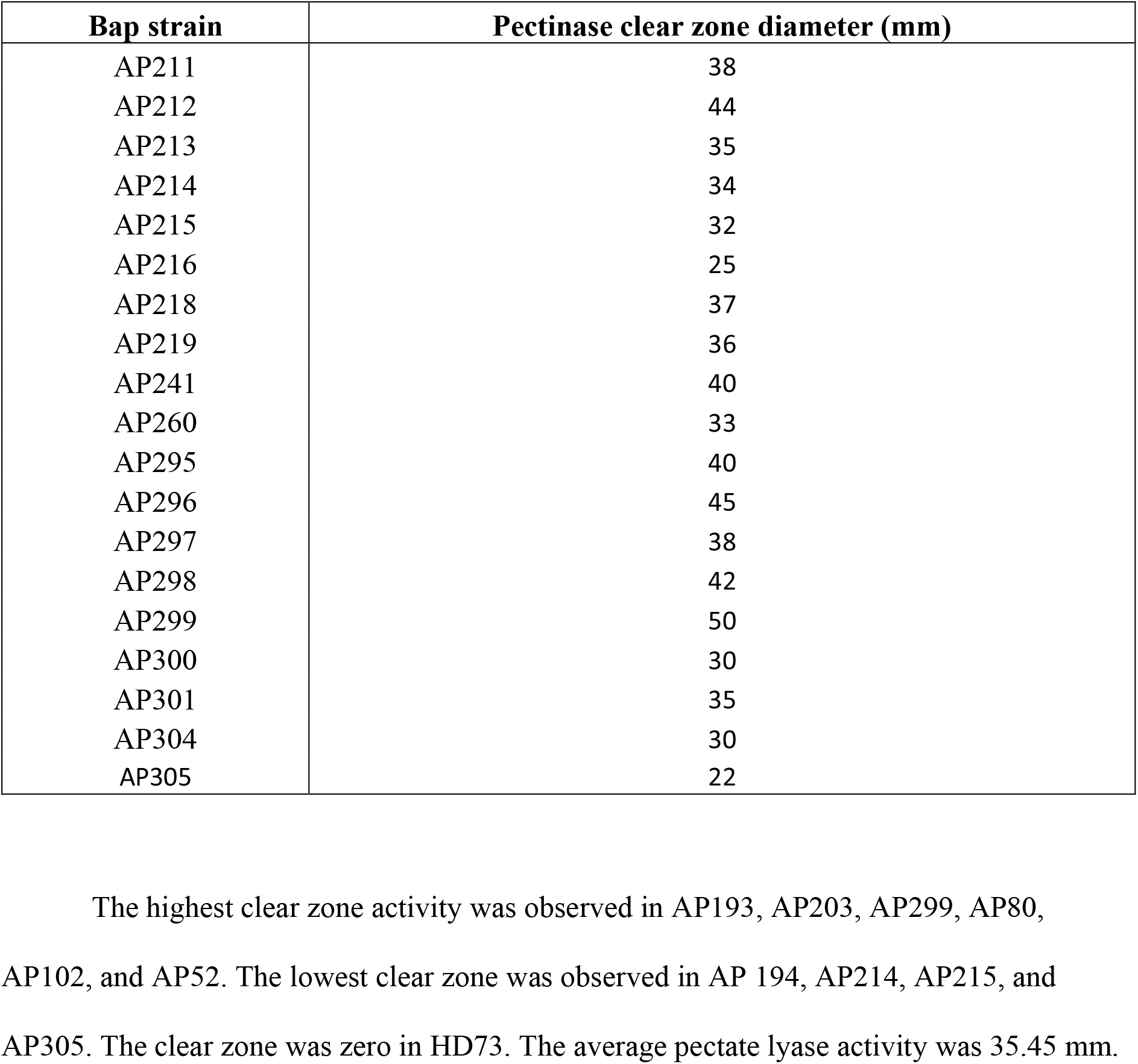
The clear zones ranges from 22 mm to 55 mm in Bap strains

### 3.3. Pectin carbon uptake test

**Table 3.**
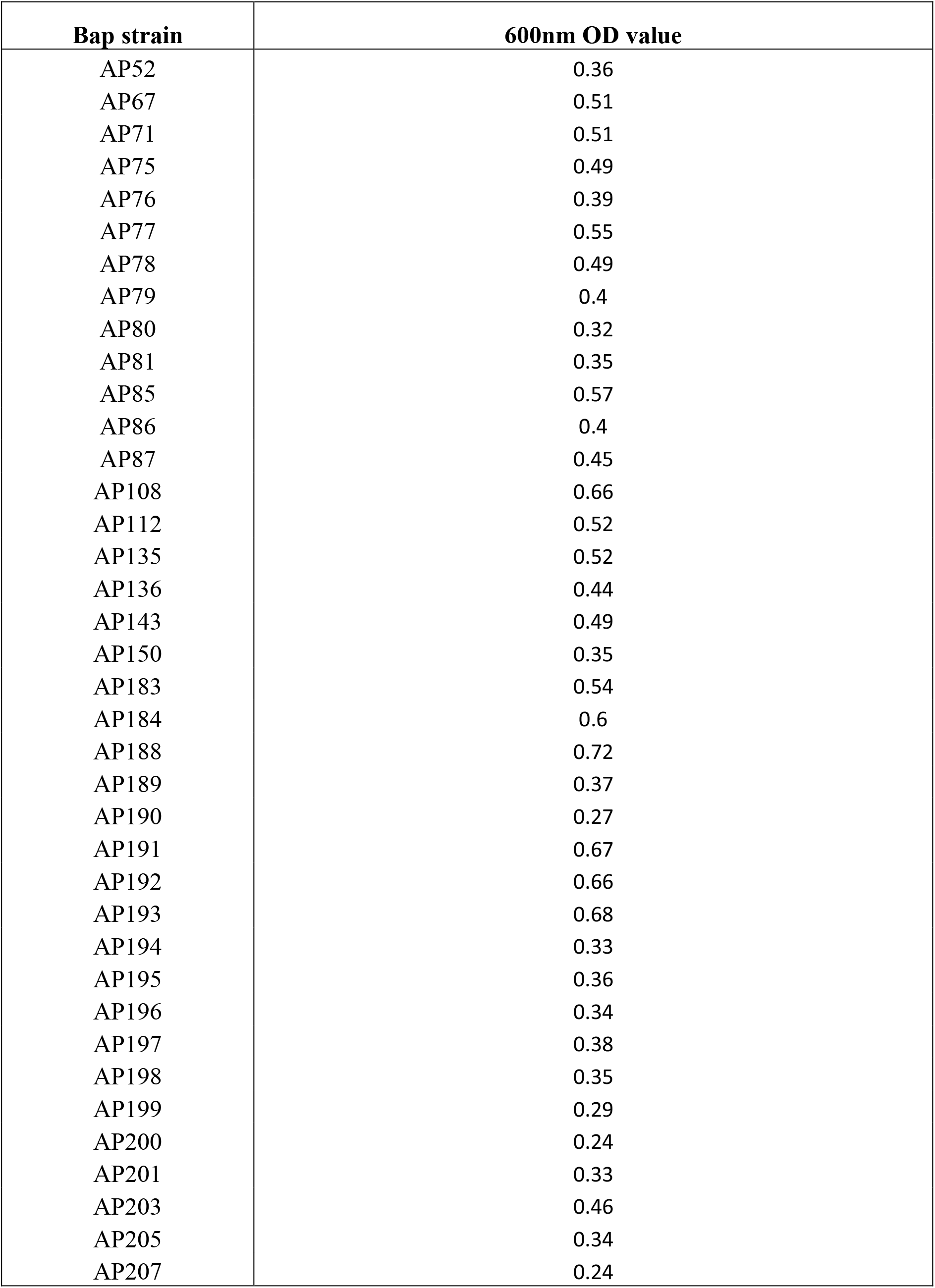
Pectin utilization of 40 Bap strains.

**Table 4.** Pectin utilization of 21 Bap strains.

| **Bap strain** | **600nm OD value** |
| --- | --- |
| AP208 | 0.38 |
| AP210 | 0.15 |
| AP211 | 0.2 |
| AP212 | 0.22 |
| AP213 | 0.29 |
| AP214 | 0.2 |
| AP215 | 0.09 |
| AP216 | 0.38 |
| AP218 | 0.1 |
| AP219 | 0.21 |
| AP241 | 0.1 |
| AP260 | 0.17 |
| AP295 | 0.18 |
| AP296 | 0.11 |
| AP297 | 0.22 |
| AP298 | 0.22 |
| AP299 | 0.19 |
| AP300 | 0.05 |
| AP301 | 0.09 |
| AP304 | 0.2 |
| AP305 | 0.11 |

**Table 5.** 12 strains showed the highest level of pectin utilization based on OD.

| **Bap strain** | **600nm OD value** |
| --- | --- |
| AP67 | 0.51 |
| AP71 | 0.51 |
| AP77 | 0.55 |
| AP78 | 0.49 |
| AP85 | 0.57 |
| AP108 | 0.66 |
| AP135 | 0.52 |
| AP143 | 0.49 |
| AP189 | 0.37 |
| AP192 | 0.66 |
| AP193 | 0.68 |

**Table 6.** Six strains showed the lowest level of pectin utilization based on OD.

| Bap strain | 600nm OD value |
| --- | --- |
| AP194 | 0.1 |
| AP 205 | 0.24 |
| AP214 | 0.09 |
| AP216 | 0.1 |
| AP218 | 0.21 |
| AP219 | 0.1 |
| HD73 | 0.09 |

### 3.4. Whole genome sequencing of Bap strains

The GC (%) contents, genome size, largest contig, sequence reads, and average coverage of AP 194 strains were 46.33, ~3.988 Mbp, 688,671bp, 1,179,872, and 28.77x. The total read count, mean read length, and total read length were 1,132,586, 100.91, and 114,293,488. The total contig length was 3,971,326 and mean contig length was 86,333. The GC (%) contents was 46.23, genome size was 4.039 Mbp, largest contig was 486,108 bp, sequence reads was 1,056,594 bp, and average coverage was 28.12x in AP 214 strains. The total read count was 1,015,508 bp, mean read length was 111.75, and total read length was 113,479,797 bp. The total contig lenth was 4,034,213 and mean contig length was 65,067.

### 3.5. RAST analysis

The number of contigs with protein coding genes was 59, the number of subsystems was 462, the number of coding sequences was 4014, and the number of RNAs was 98 in AP194 bap strain. The number of contigs with protein coding genes was 51, the number of subsystems was 462, the number of coding sequences was 4060, and the number of RNAs was 75 in AP214 bap strain. A total of 14 and 13 pectin-associated subsystem feature were found in AP194 and AP214. The altronate hydrolase (uxaA) gene was found in AP194 Bap strain. The uxaB gene was not found in AP214 Bap strain.

### 3.6. CLC genomics analysis of exuT and uxuB protein gene sequence alignment

In translation frame three of *exuT* gene sequence alignment, glycine (G) amino acid was found in AP194 and AP214 Bap strains. However, cysteine (C) amino acid was found in the same translation frame of reference strains CC178, FZB42, Trigocor1448, CAU-B946, IT-45, and AS43.3. cysteine (C) also was available in AP193 Bap strains. Isoleucine (I) was found only in AP214 Bap strains. phenylalanine (F) was found in AP193, AP194, and other reference strains.

In translation frame one and two of *exuT* gene sequence alignment, arginine (R) amino acid was found in AP194 and AP214 Bap strains. On the contrary, glycine (G), and leucine (L) amino acid were found in the same translation frame of reference strains CC178, FZB42, Trigocor1448, CAU-B946, IT-45, and AS43.3. Arginine (R) also was found in AP193 Bap strains.

In translation frame two and three of *uxuB* gene sequence alignment, isoleucine (I), tryptophan (W), methionine (M), tyrosine (Y), and histidine (H) were found in AP 194, and AP214 Bap strains. However, threonine (T), cysteine (C), valine (V), leucine (L), and arginine (R) amino acid were found in the same translation frame of reference strains UCMB5113, UCMB5033, CC178, FZB42, Trigocor1448, CAU-B946, IT-45, NAU-B3, Y2, and AS43.3.

## 4. Discussion

The presented results indicate that Bap strains had slower growth rates on pectate agar media than on TSA media. From a total of 59 Bap strains, three strains (AP207, AP214, and AP215) did not grow very well on PA media. The remaining 56 Bap strains growth had started after 6 hr inoculation time. The pectate lyase test results demonstrate that six Bap strains had the largest clear zone around the bacterial colony. Two Bap strains (AP194 and AP214), and one strain (HD73) had the lowest and zero clear zone around the colony. Based on the results, it can be concluded that the highest clear zone showing strains have the highest pectate lyase activity. In contrast, the lowest clear zone showing strains had the lowest pectate lyase activity, and zero clear zone strain has no activity. Previous studies found that the clear zone formed in *Bacillus sp.* Strain KSM-P15 (Kobayashi et al., 1999) in 10 minutes. However, the clear zone around the Bap strains colony were observed after 30 minutes.

Based on the pectin carbon uptake test, 12 *Bap* strains grew vigorously in 1% pectin. The remaining *Bap* strains have showed lowest and average growth in TSS medium. Two *Bap* strains AP194 and AP214 did not grow in TSS medium. *Btk* strain HD73 also did not grow in TSS medium. Because, *Btk* strain HD73 had no pectin-associated genes. It can be inferred from the results that *exuT* and *uxuB* pectin-associated genes from the fastest growing strains might transport and metabolize faster than other strains.

Rast results of AP193, AP194, and AP214 Bap strains have revealed that they have differences among them. Pectin pathways associated subsystem features have counted in three Bap strains are 16, 14, and 13. Altronate hydrolase *(uxaB)* gene has found in AP193 and AP194 only, not present in AP214 Bap strain. However, pairwise Blastx with different reference genome sequences has indicated that it is present in three Bap gene sequences. It is also found that pectin-associated rest of the genes are present in three Bap strains.

Based on the *exuT* protein gene sequence alignment results (figure 17), it can be concluded that Glycine (G), and Arginine (R) amino acid differences exist in Bap strains AP194 and AP214 in comparison with reference strain FZB42 (Chen et al., 2007). This type of differences might affect the hexuronate transport of D-glucuronate and D-galacturonate chemical compounds into the bacterial cell.

The *uxuB* protein gene sequence alignment results (figure 16) indicate that Isoleucine (I), Tryptophan (W), Methionine (M), Tyrosine (Y), and Histidine (H) amino acid changes occurred in AP194 and AP214 in comparison with reference strain FZB42 (Chen et al., 2007). This change could hamper the overall metabolic activity of strains AP194 and AP214.

## *In vitro* pectin utilization tests on Bap strains

**Figure 1:**
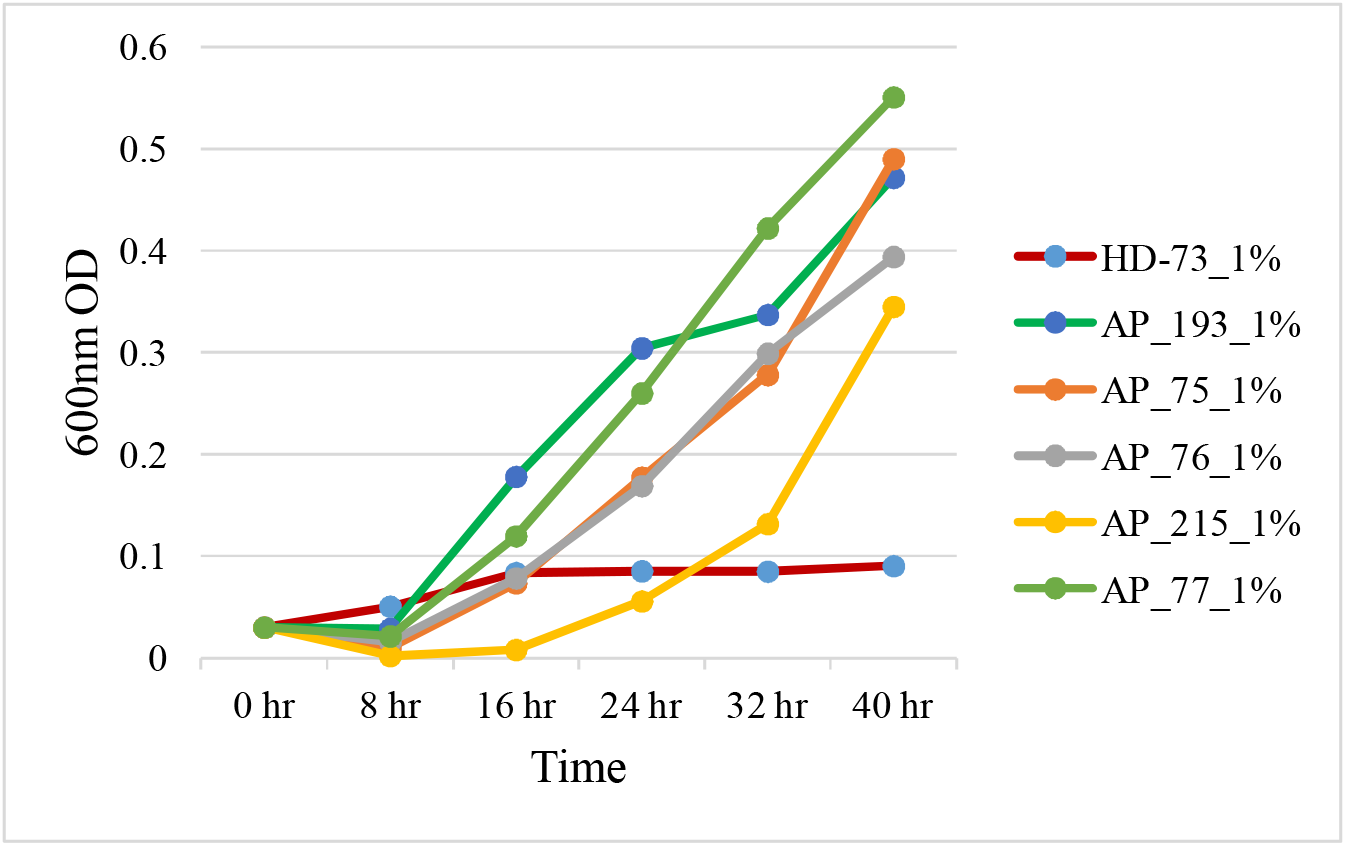
*in vitro* Bap bacterial growth using pectin as a sole carbon source (HD73, AP193, AP75, AP76, AP77, and AP215).

**Figure 2:**
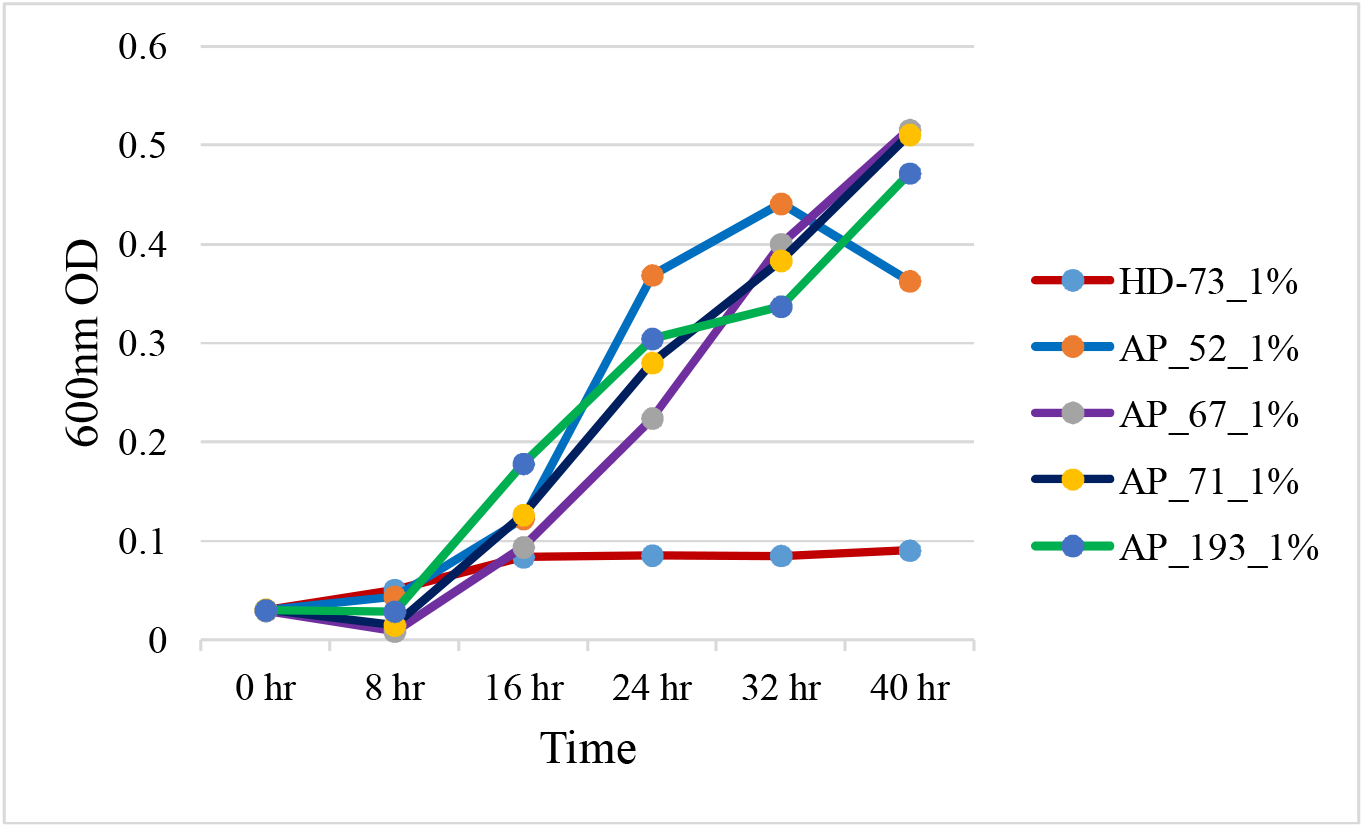
*in vitro* Bap bacterial growth using pectin as a sole carbon source (HD73, AP52, AP67, AP71, and AP193).

**Figure 3:**
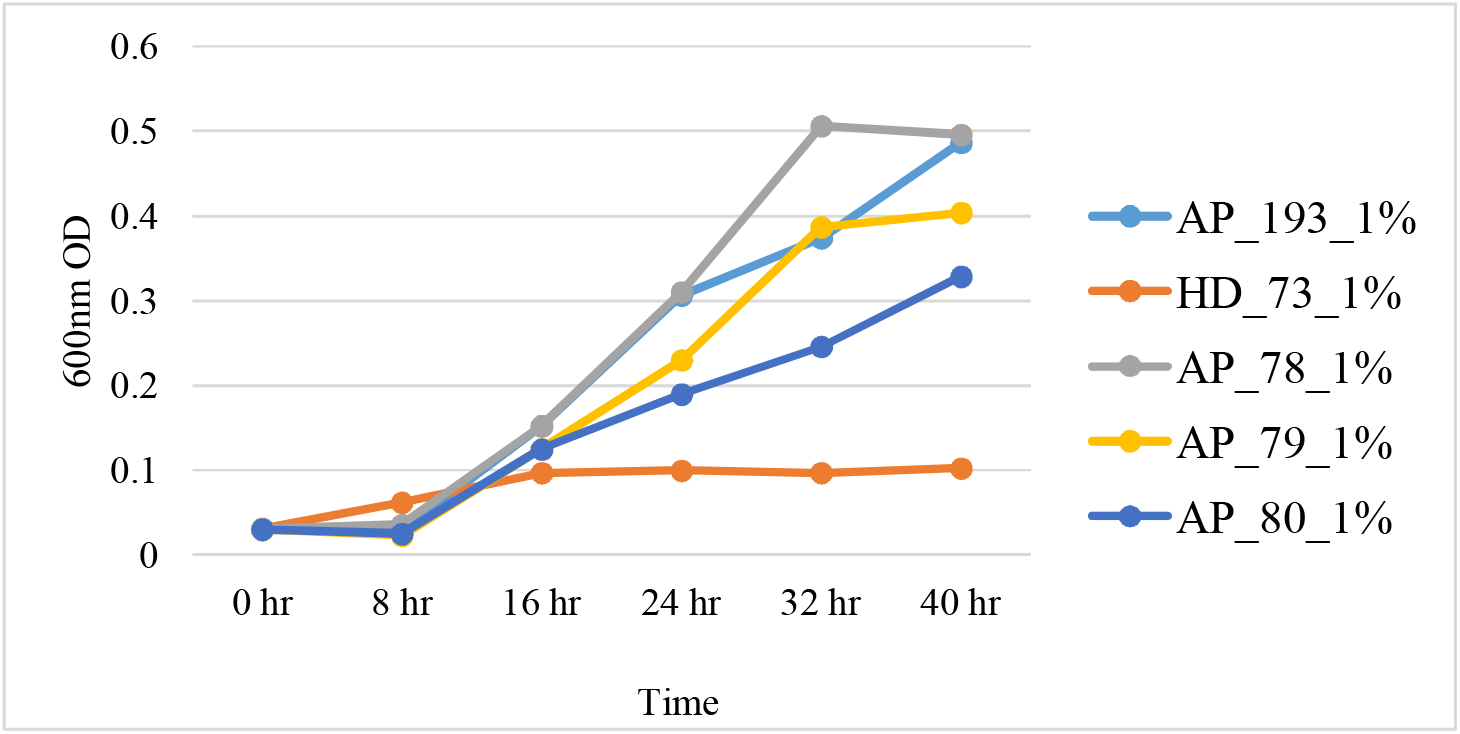
*in vitro* Bap bacterial growth using pectin as a sole carbon source (HD73, AP193, AP73, AP78, AP79, and AP80).

**Figure 4:**
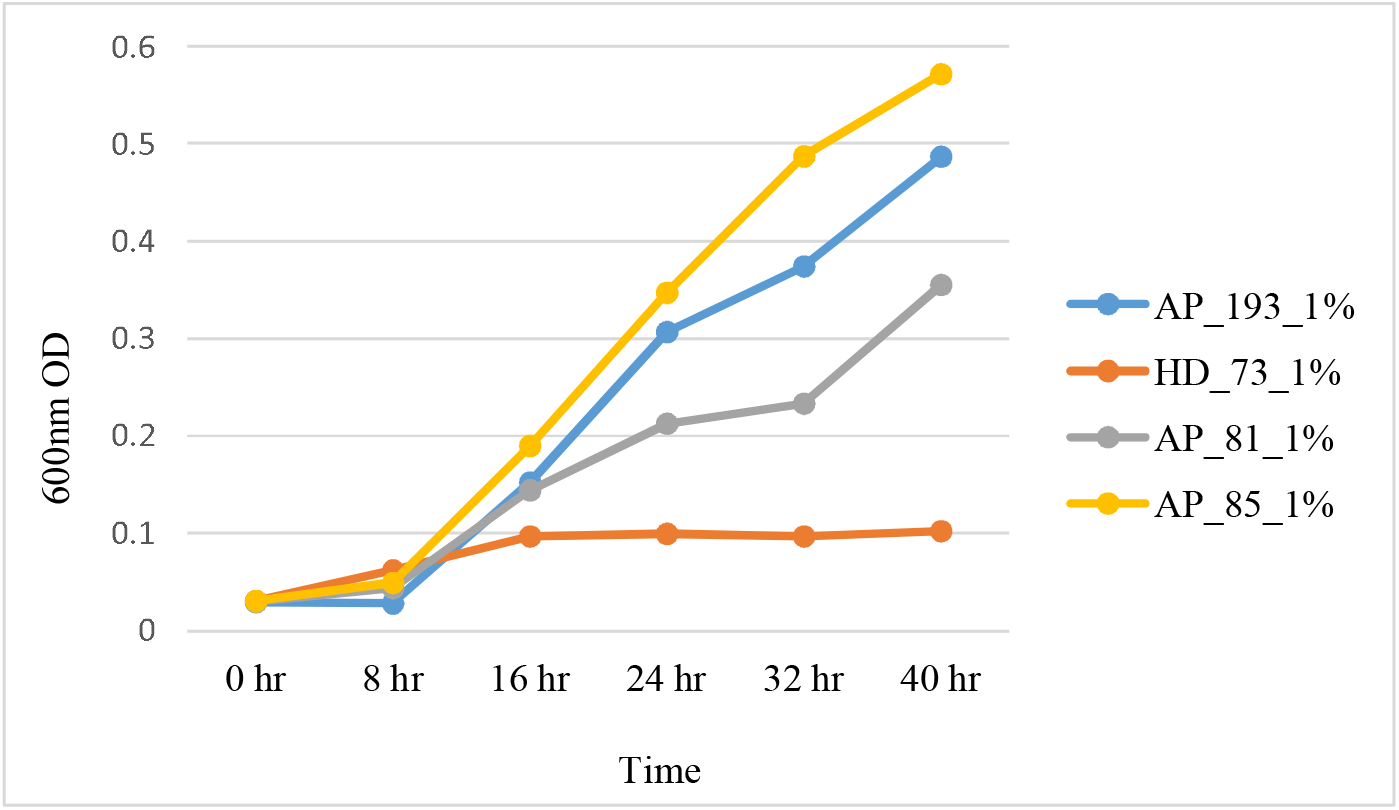
*in vitro* Bap bacterial growth using pectin as a sole carbon source (HD73, AP193, AP81, and AP85).

**Figure 5:**
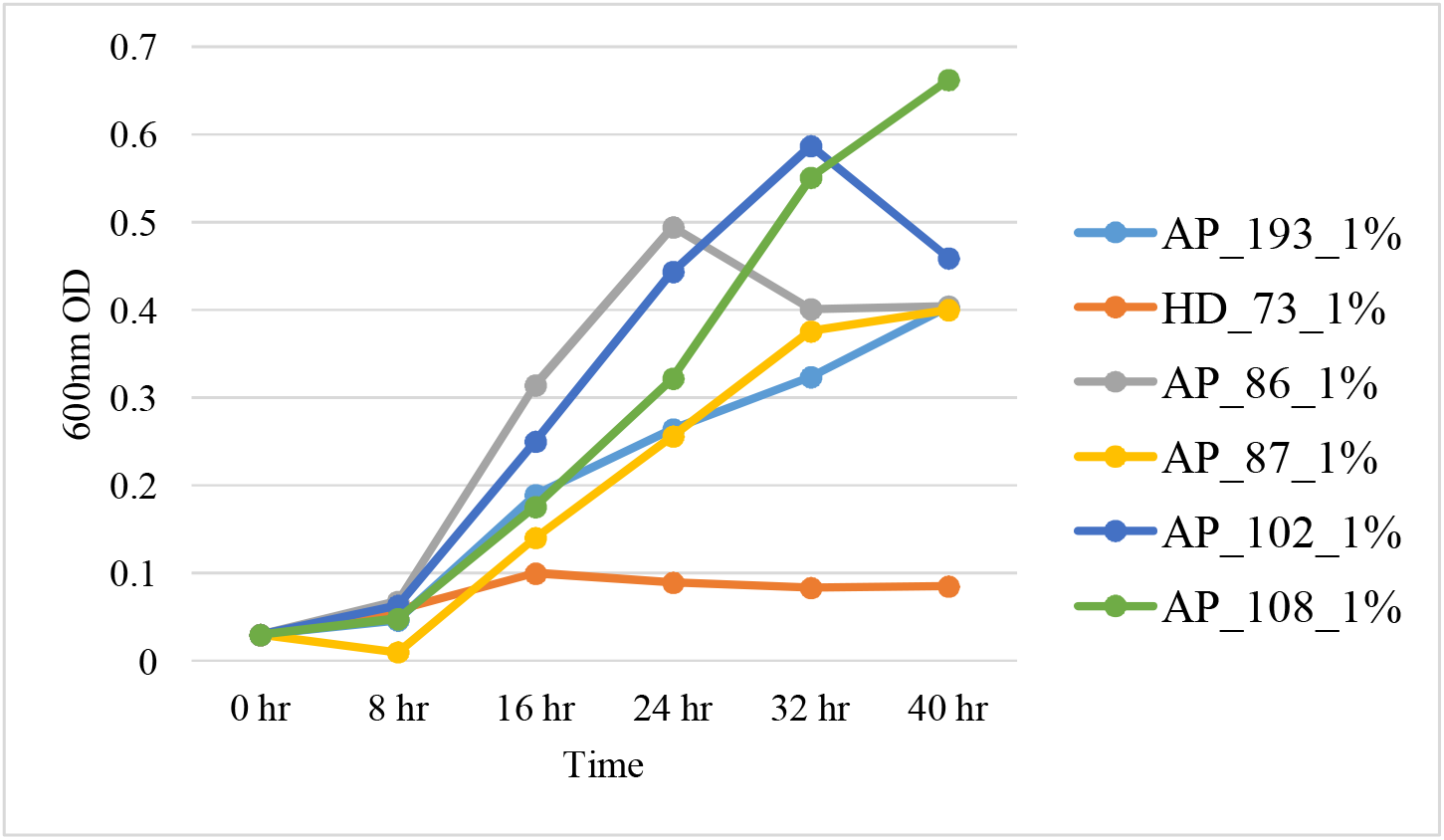
*in vitro* Bap bacterial growth using pectin as a sole carbon source (HD73, AP193, AP86, AP87, AP102, and AP108).

**Figure 6:**
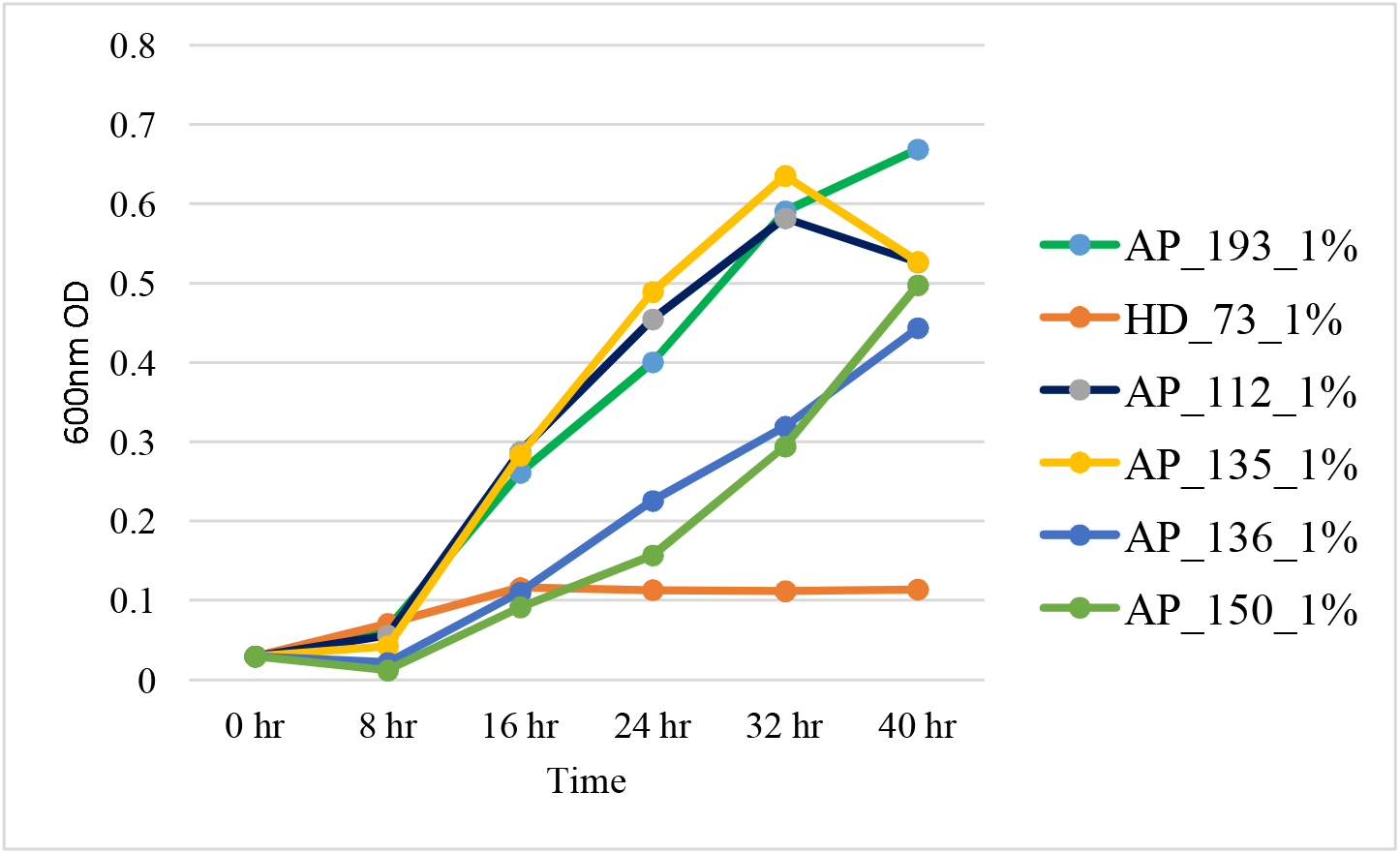
*in vitro* Bap bacterial growth using pectin as a sole carbon source (HD73, AP193, AP112, AP135, AP136, and AP150).

**Figure 7:**
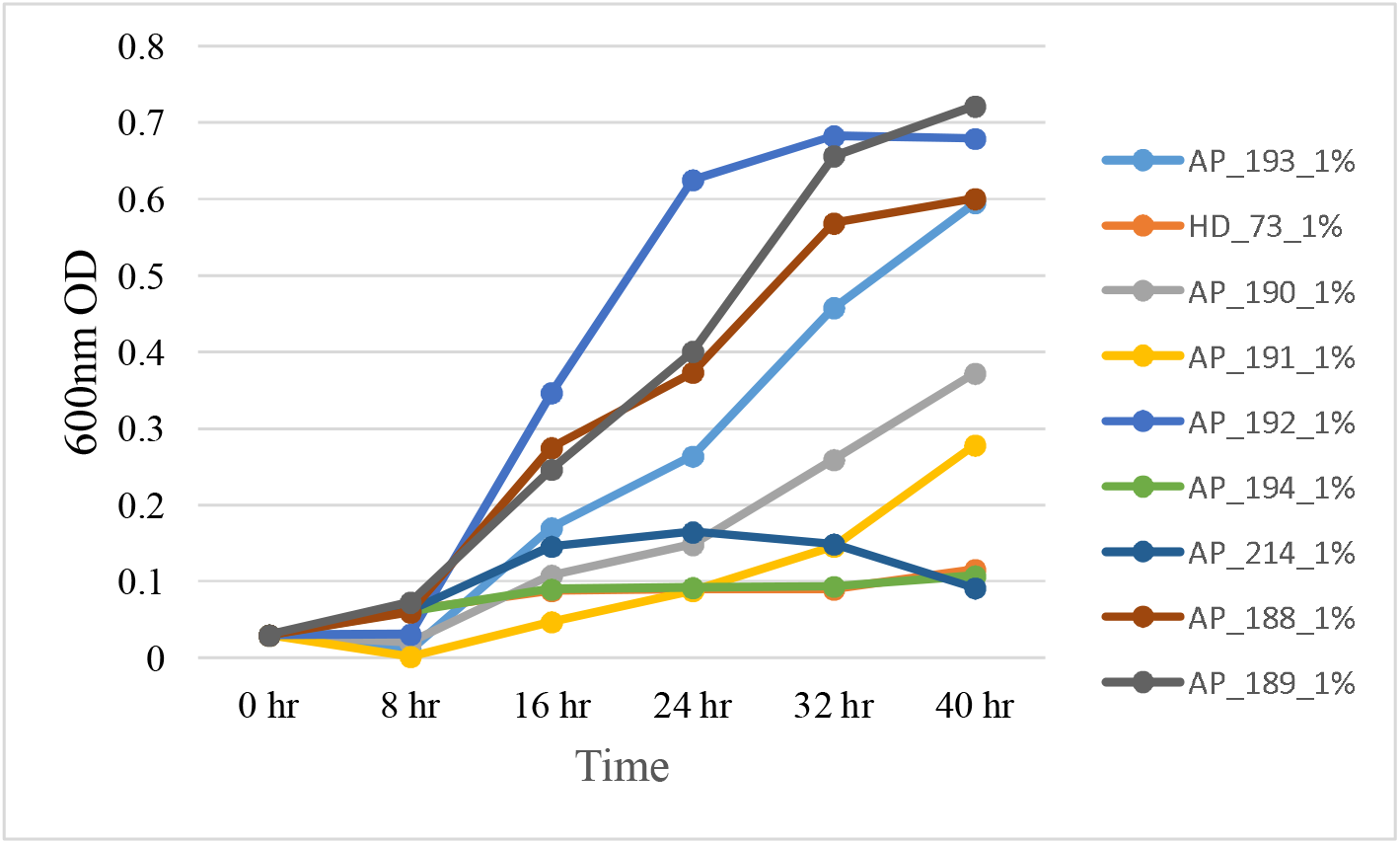
*in vitro* Bap bacterial growth using pectin as a sole carbon source (HD73, AP193, AP190, AP191, AP192, AP194, AP214, AP188, and AP189).

**Figure 8:**
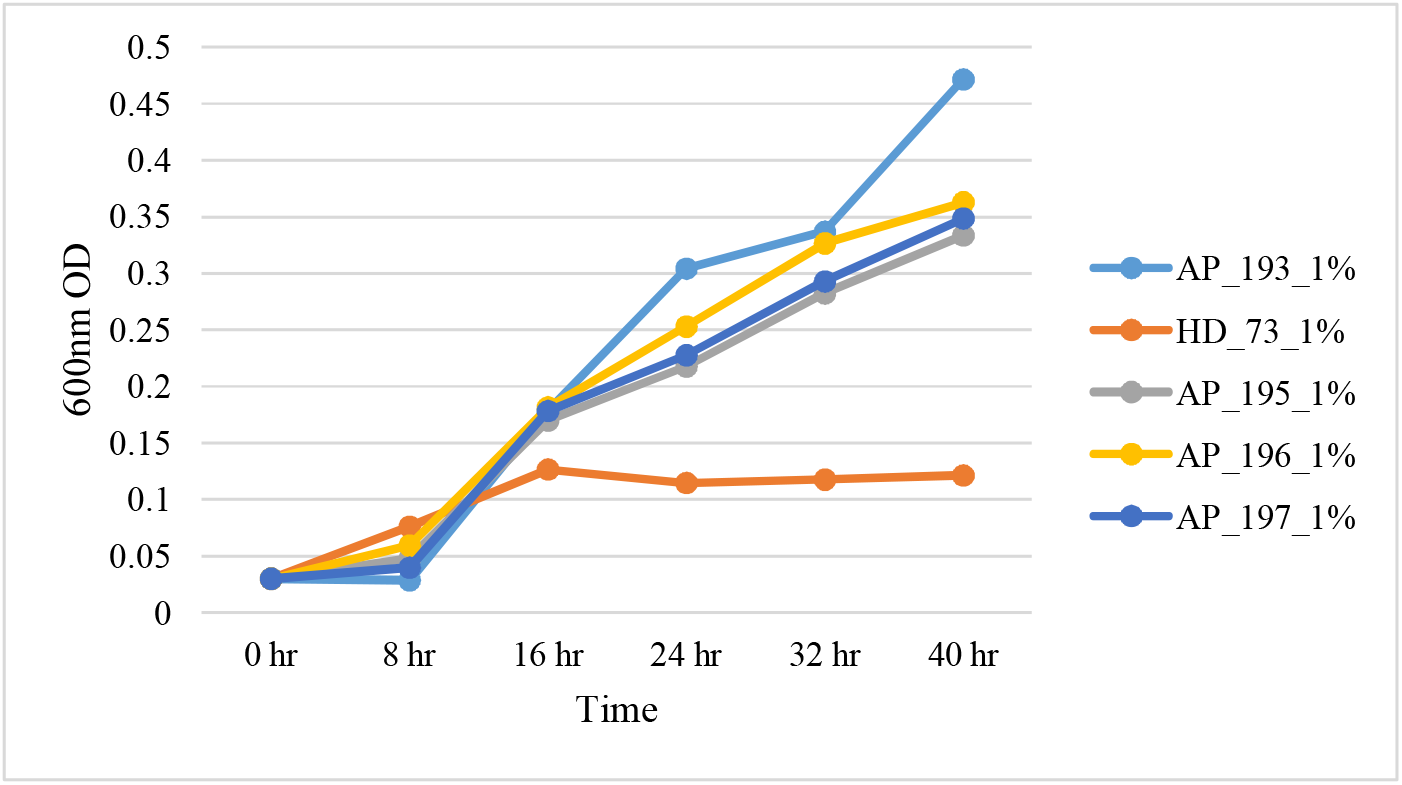
*in vitro* Bap bacterial growth using pectin as a sole carbon source (HD73, AP193, AP195, AP196, and AP197).

**Figure 9:**
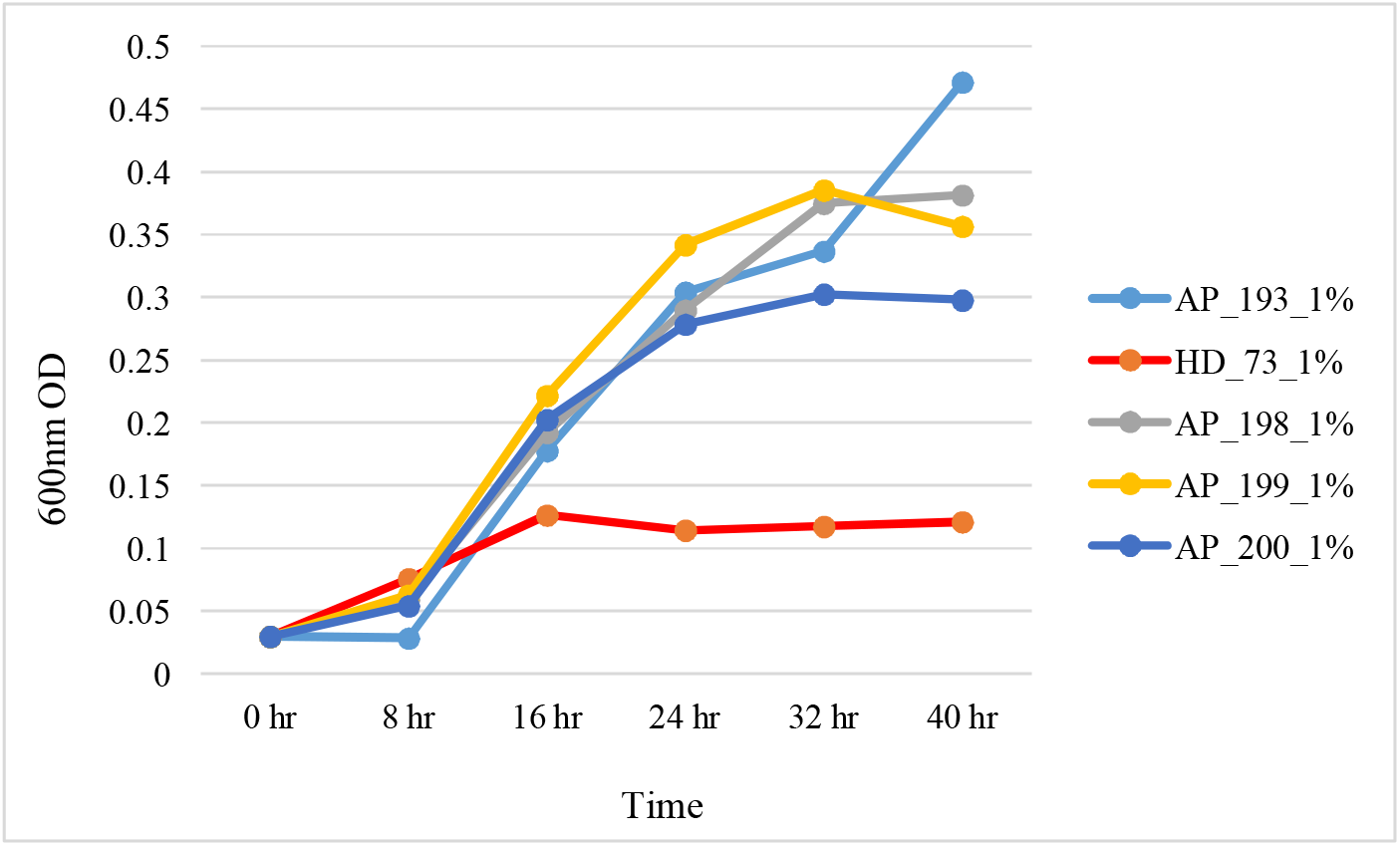
*in vitro* Bap bacterial growth using pectin as a sole carbon source (HD73, AP193, AP198, AP199, and AP200).

**Figure 10:**
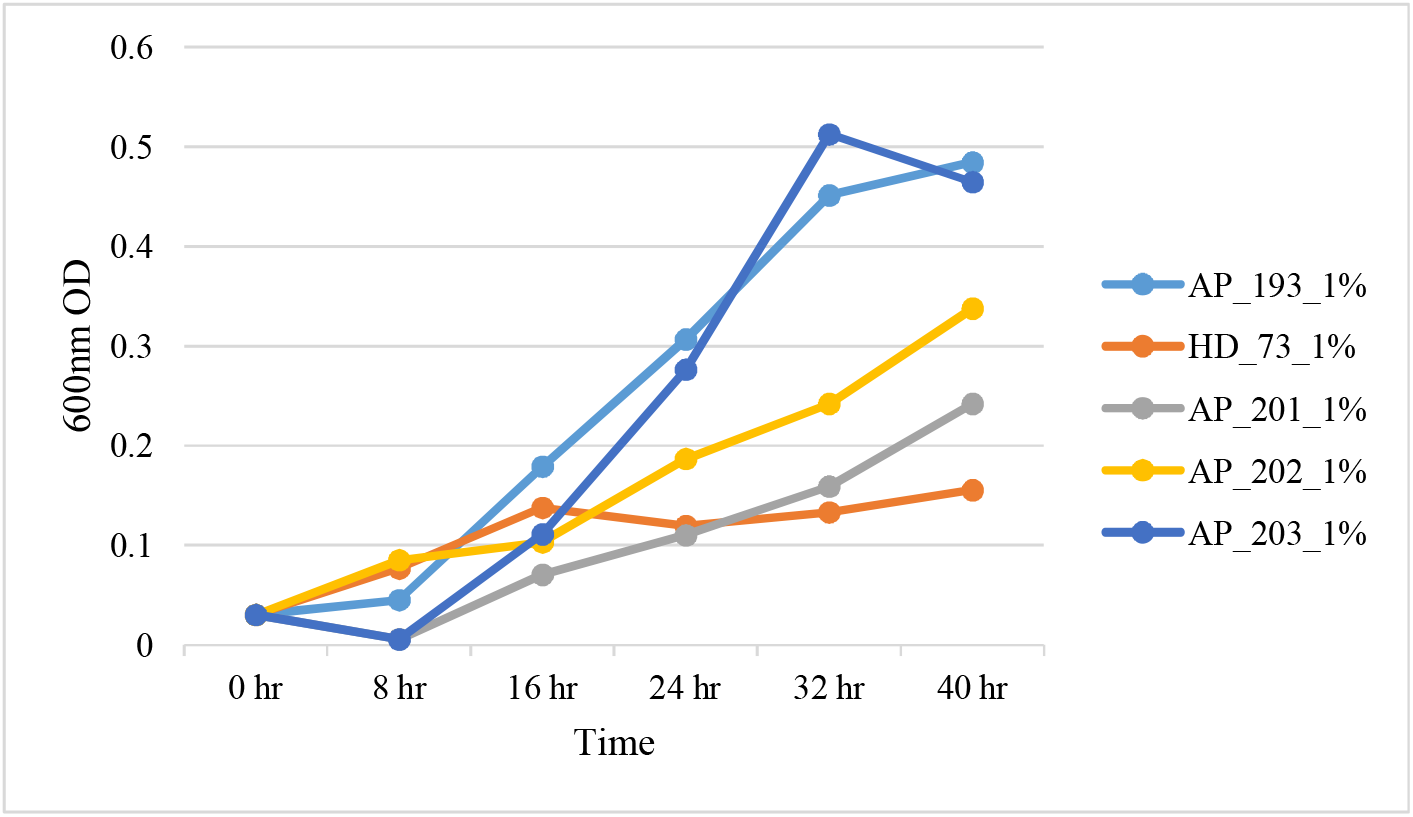
*in vitro* Bap bacterial growth using pectin as a sole carbon source (HD73, AP193, AP201, AP202, and AP203).

**Figure 11:**
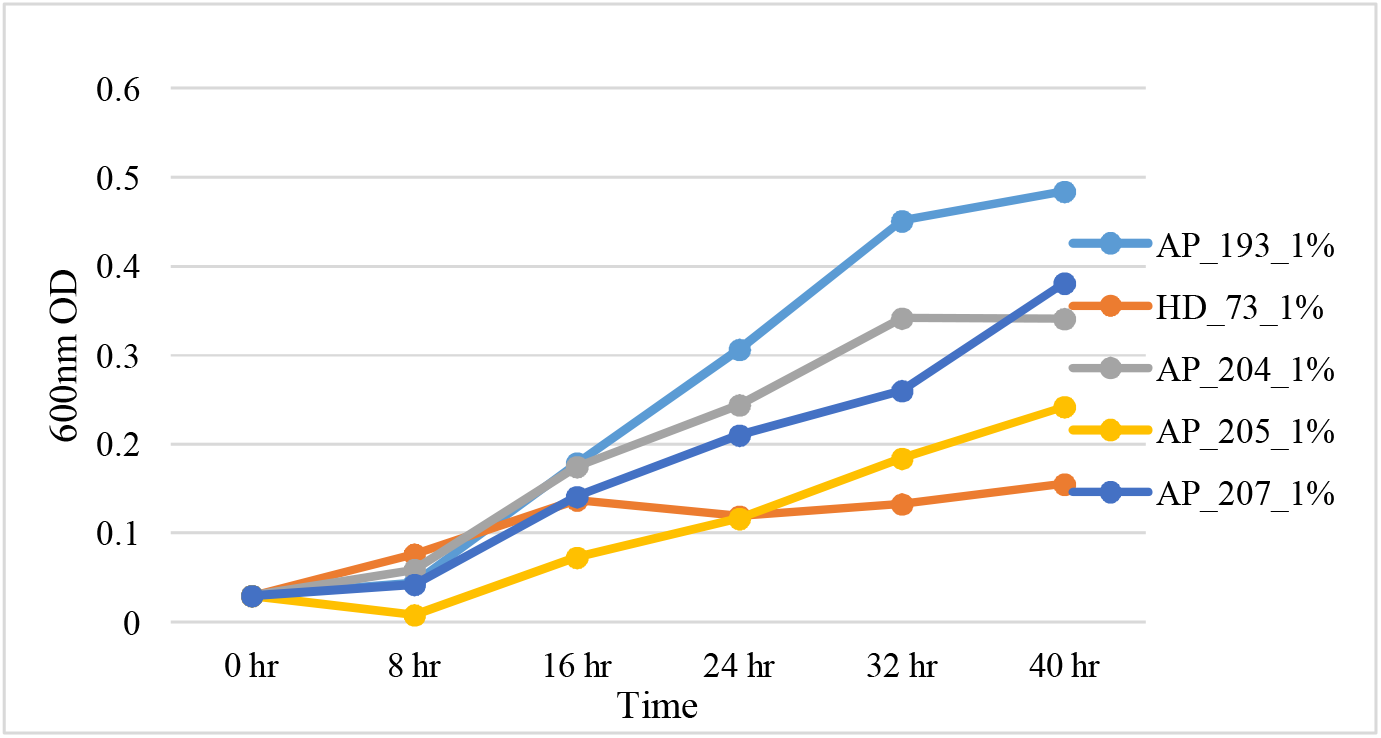
*in vitro* Bap bacterial growth using pectin as a sole carbon source (HD73, AP193, AP204, AP205, and AP207).

**Figure 12:**
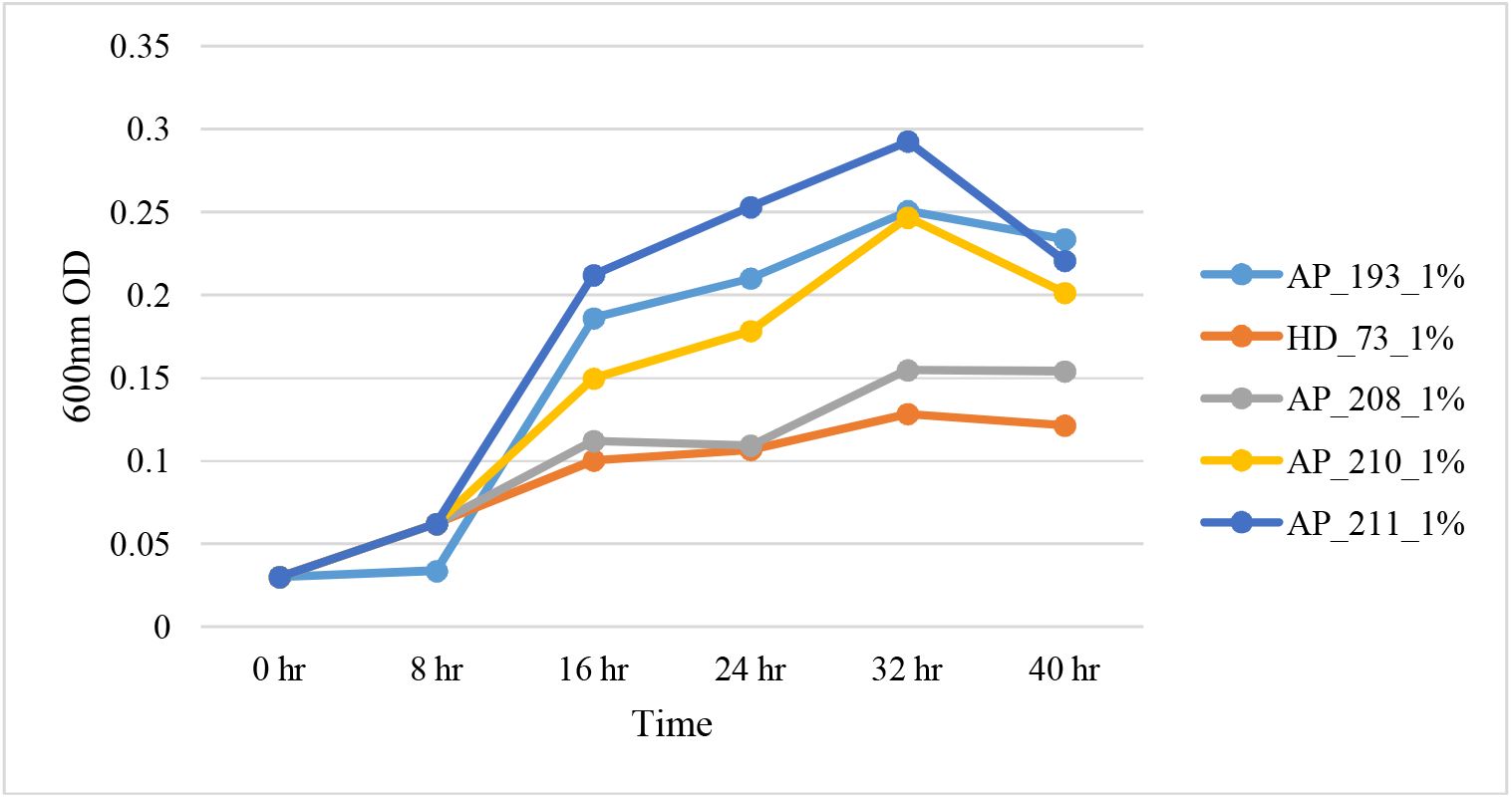
*in vitro* Bap bacterial growth using pectin as a sole carbon source (HD73, AP193, AP208, AP210, and AP211).

**Figure 13:**
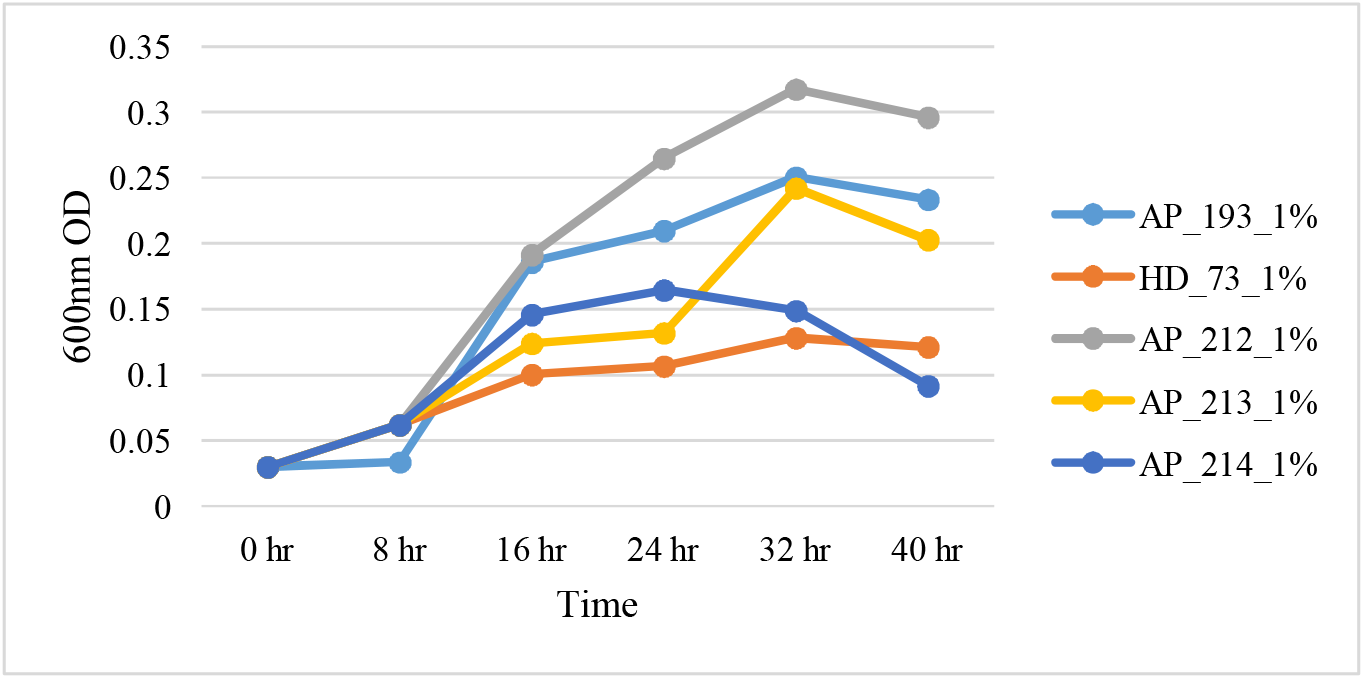
*in vitro* Bap bacterial growth using pectin as a sole carbon source (HD73, AP193, AP212, AP213, and AP214).

**Figure 14:**
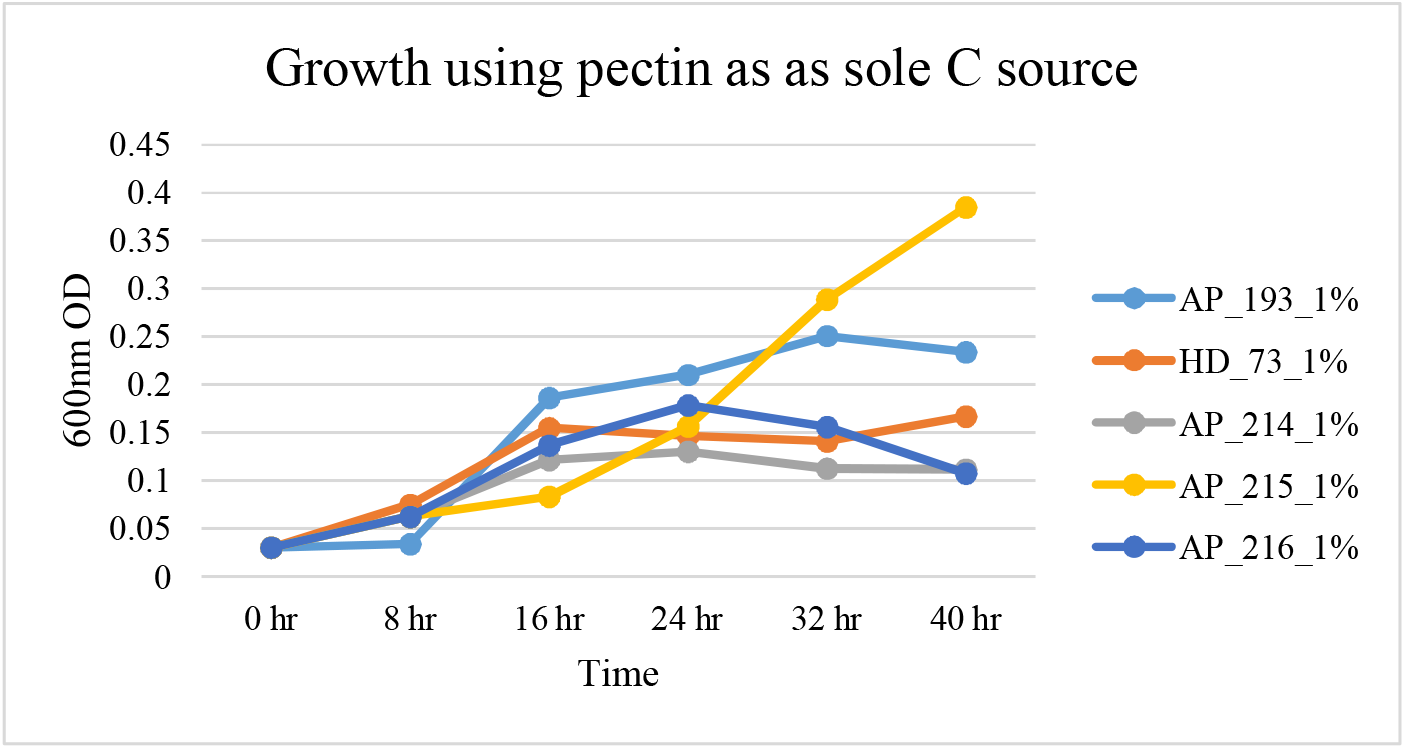
*in vitro* Bap bacterial growth using pectin as a sole carbon source (HD73, AP193, AP214, AP215, and AP216).

**Figure 15:**
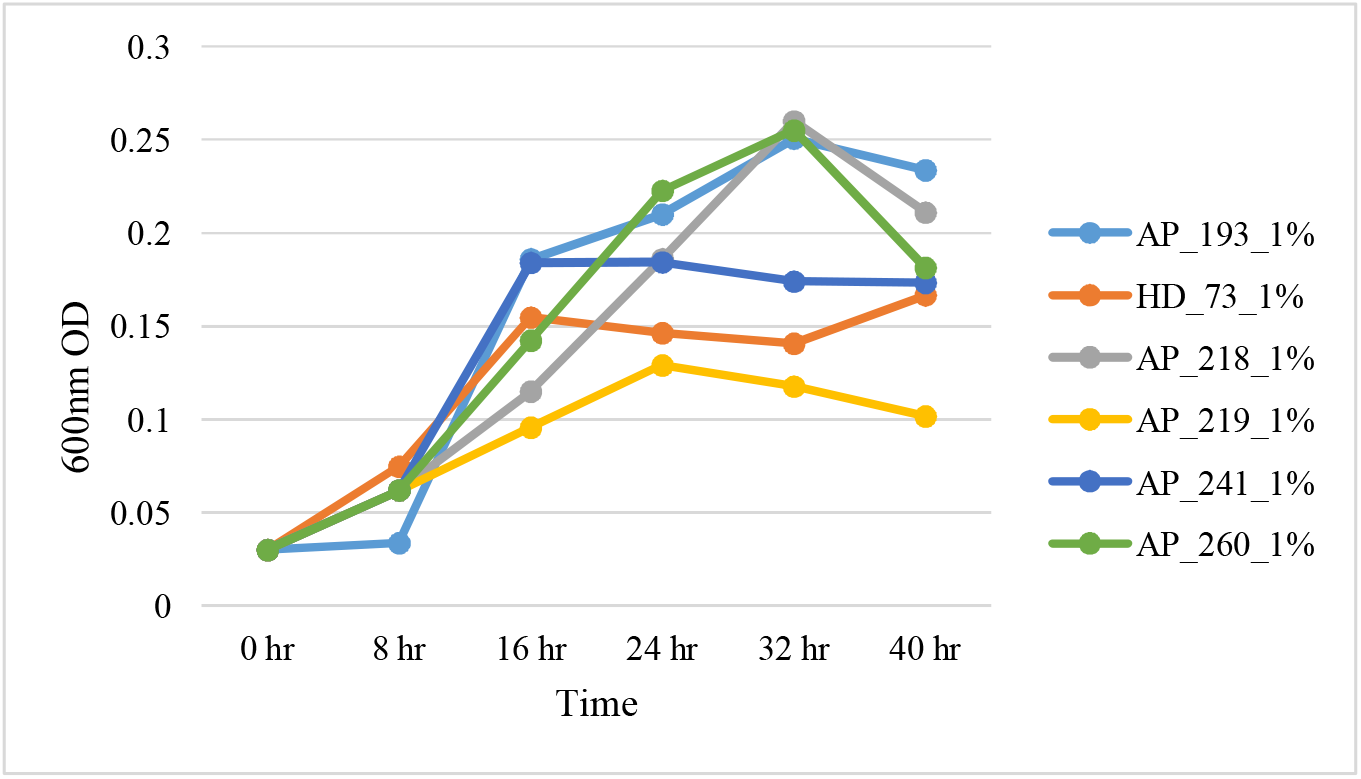
*in vitro* Bap bacterial growth using pectin as a sole carbon source (HD73, AP193, AP218, AP219, AP241, and AP260).

**Figure 16:**
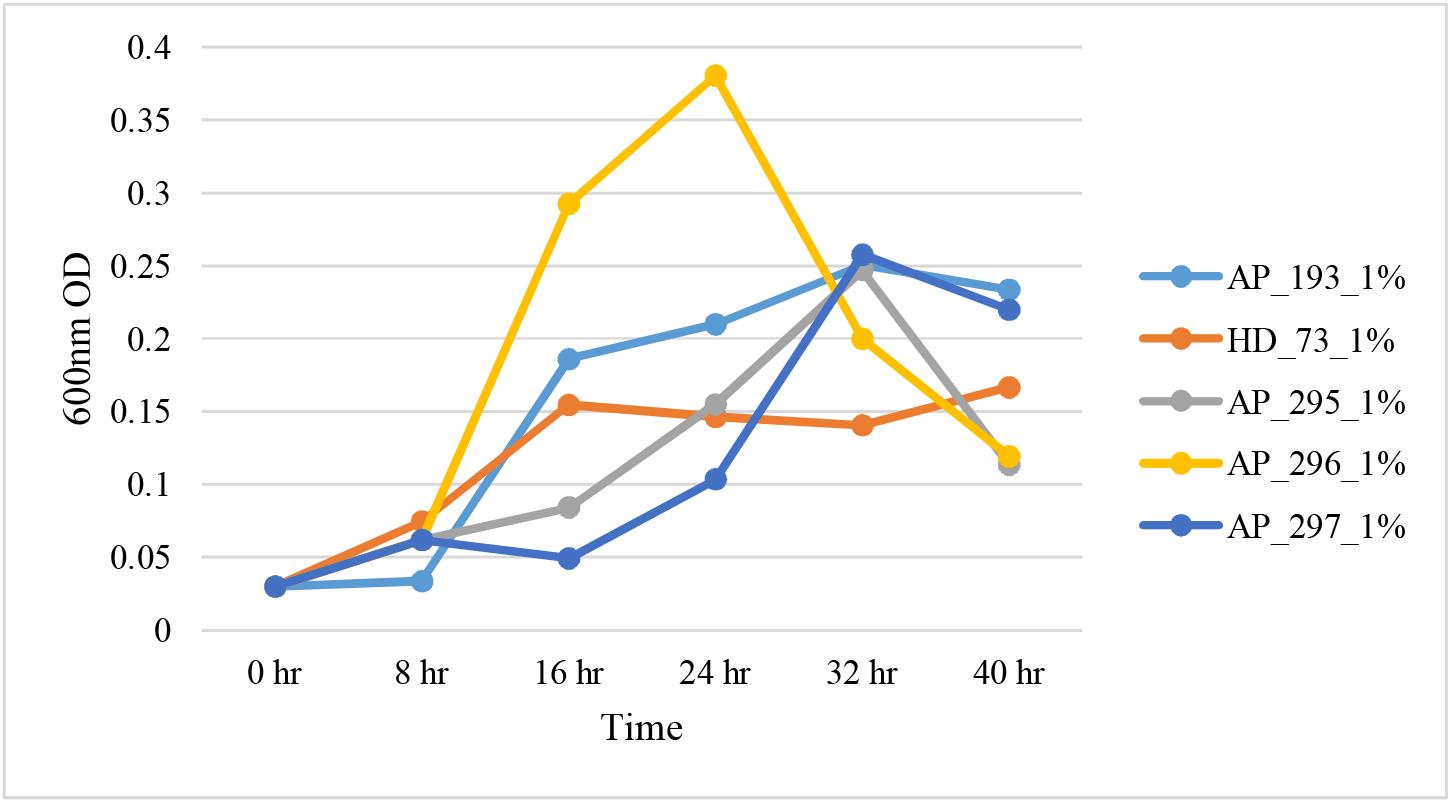
*in vitro* Bap bacterial growth using pectin as a sole carbon source (HD73, AP193, AP295, AP296, and AP297).

**Figure 17:**
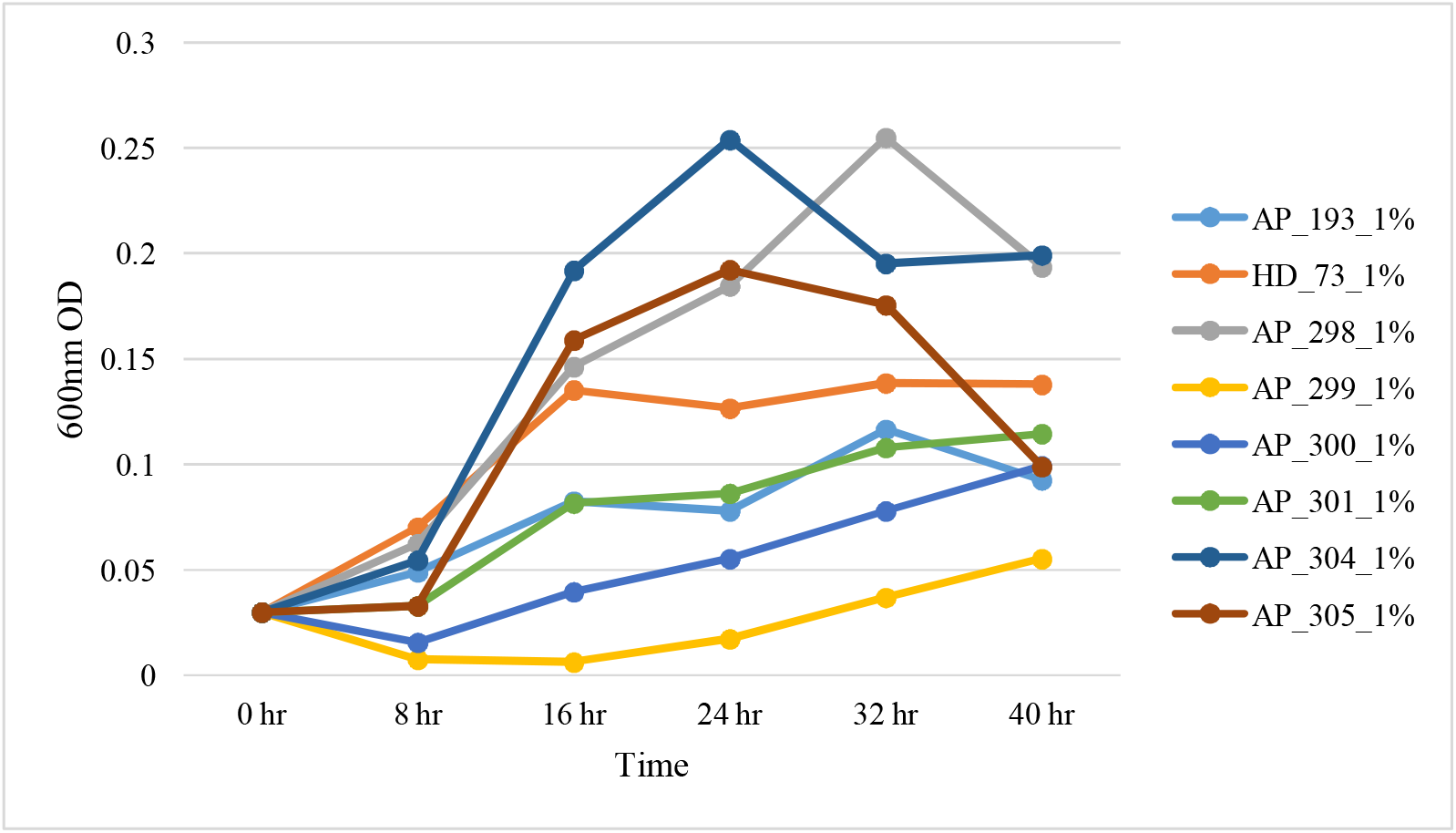
*in vitro* Bap bacterial growth using pectin as a sole carbon source (HD73, AP193, AP298, AP299, AP300, AP301, AP304, and AP305).

## Pectate lyase activity of Bap strains

**Figure 18:**
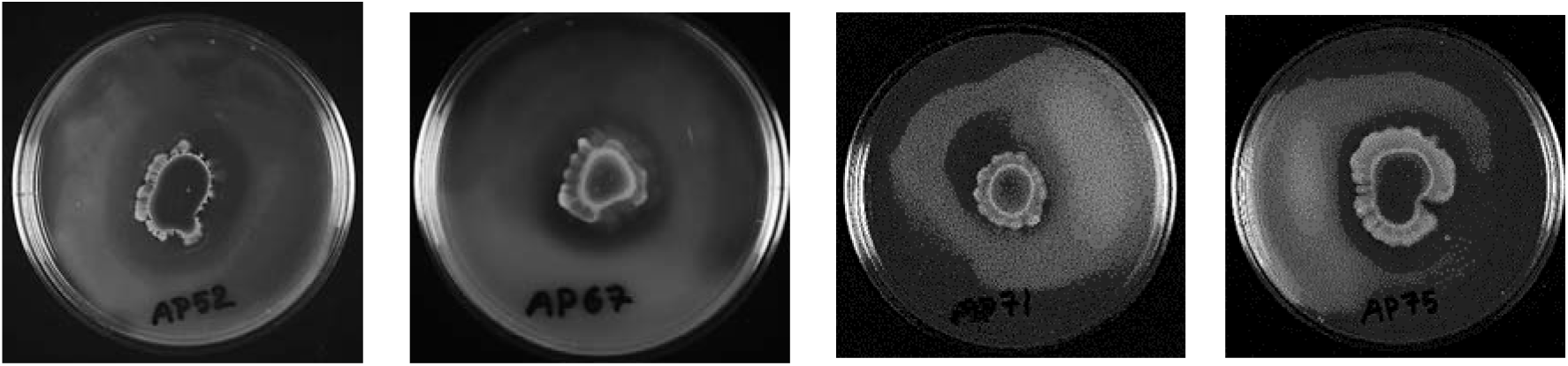
Pectate lyase activity of Bap strains (AP52, AP67, AP71, and AP75).

**Figure 19:**
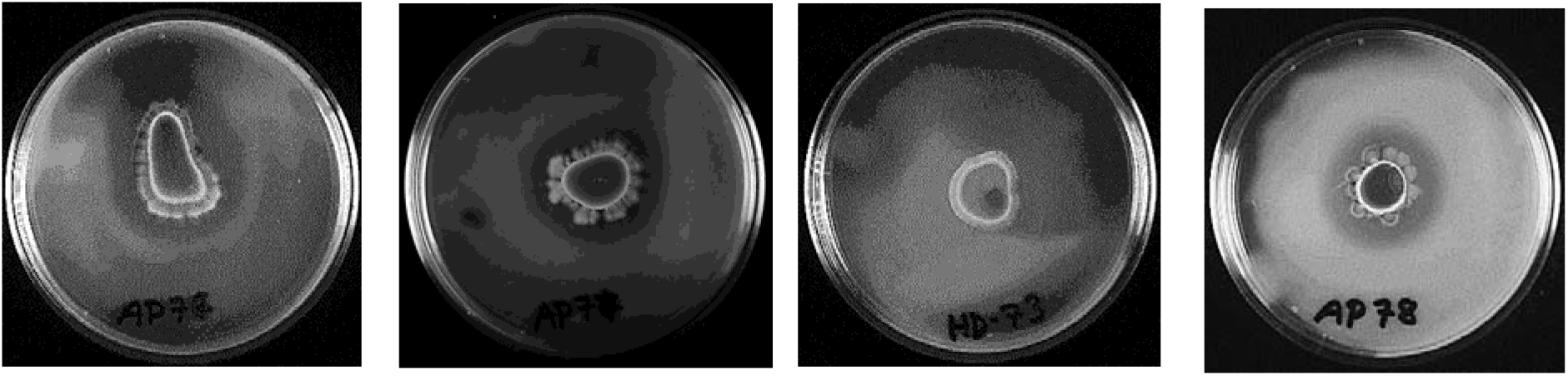
Pectate lyase activity of Bap strains (AP76, AP73, HD73, and AP78).

**Figure 20:**
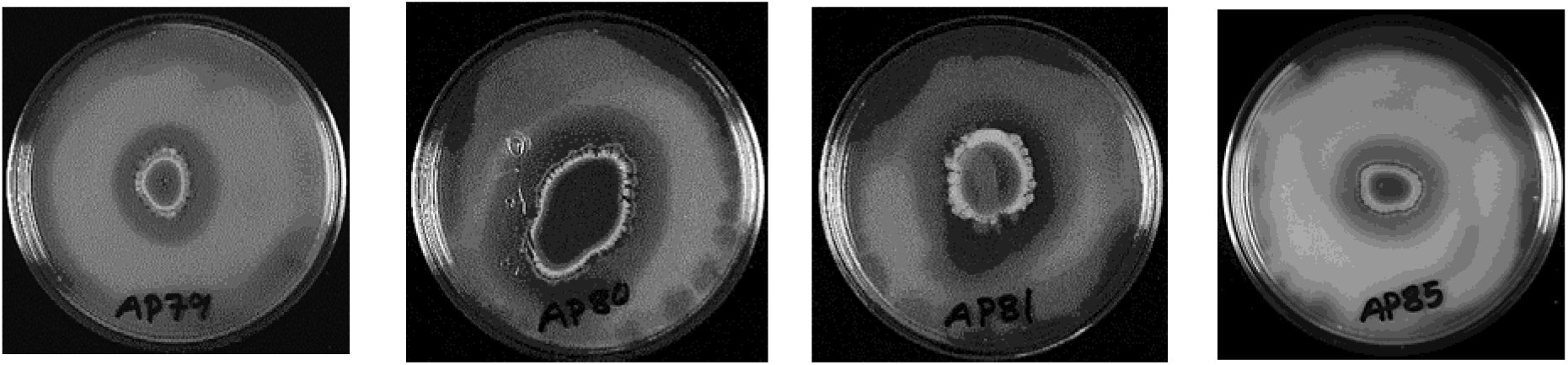
Pectate lyase activity of Bap strains (AP79, AP80, AP81, and AP85).

**Figure 21:**
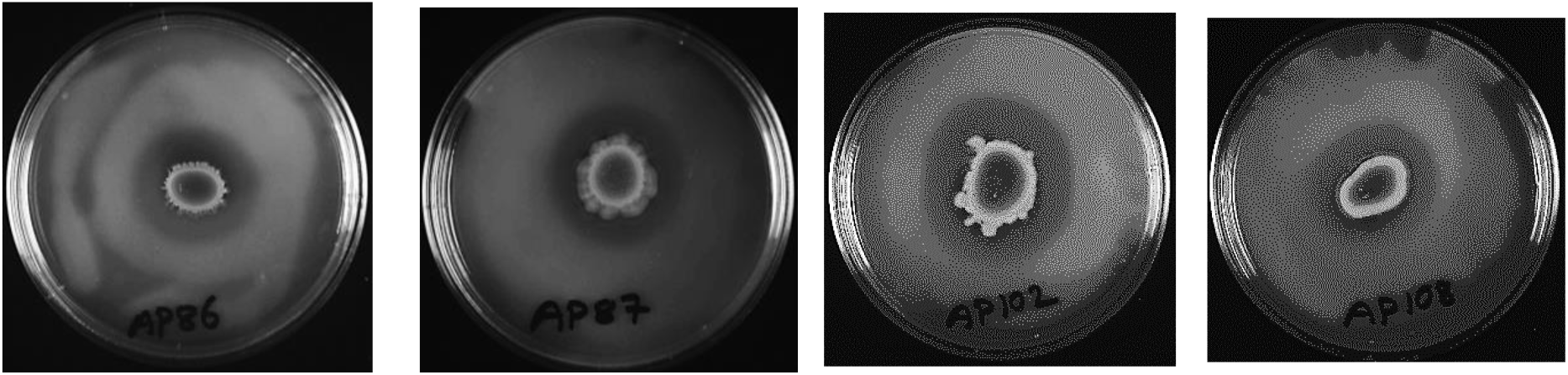
Pectate lyase activity of Bap strains (AP86, AP87, AP102, and AP108).

**Figure 22:**
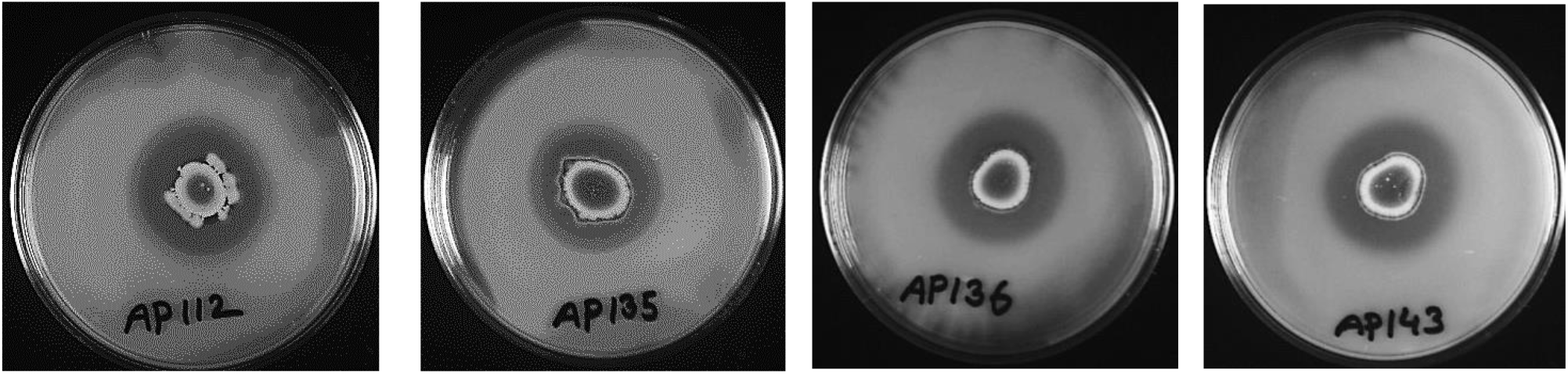
Pectate lyase activity of Bap strains (AP112, AP135, AP136, and AP143).

**Figure 23:**
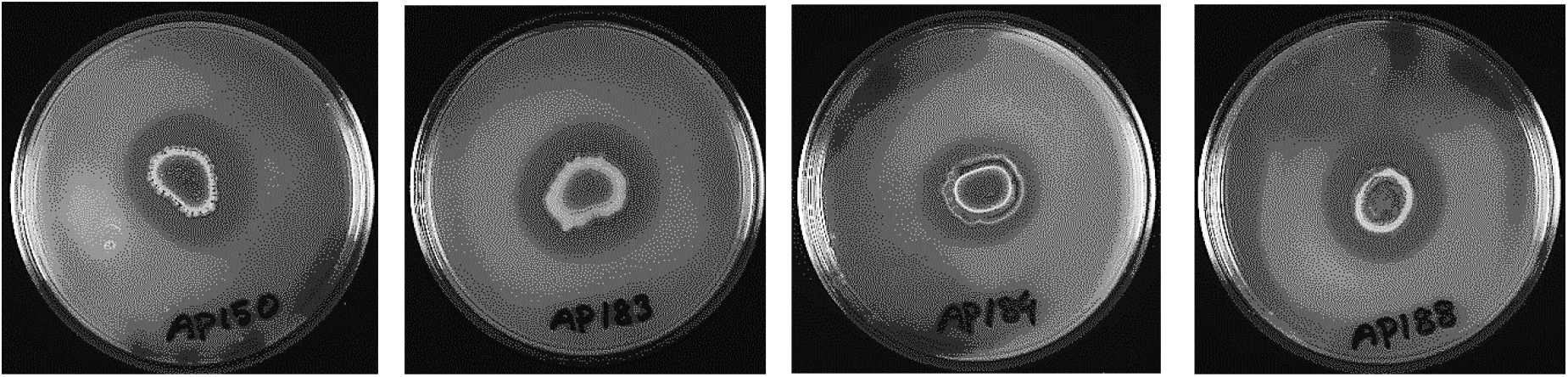
Pectate lyase activity of Bap strains (AP180, AP183, AP184, and AP188).

**Figure 24:**
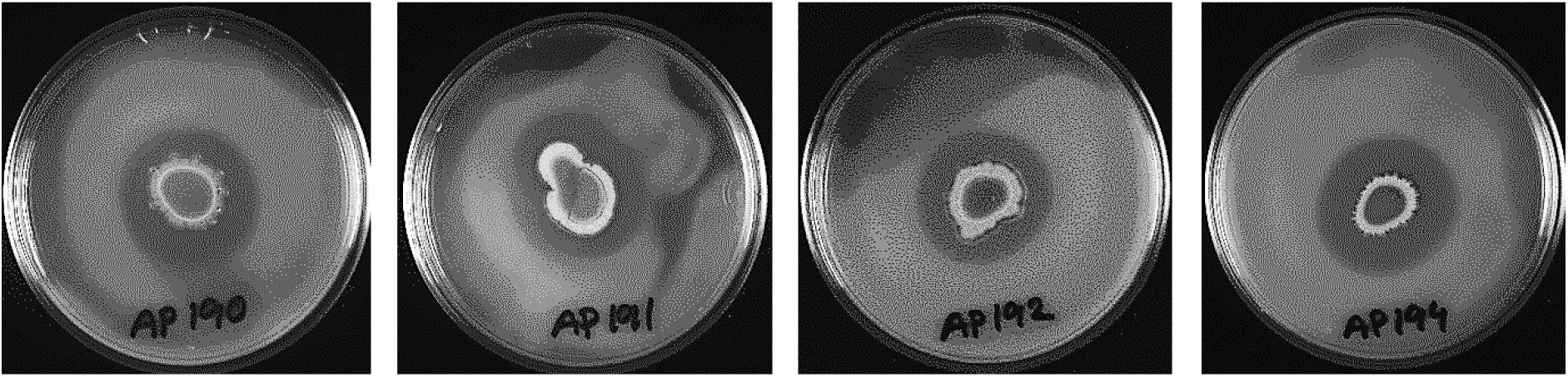
Pectate lyase activity of Bap strains (AP190, AP191, AP192, and AP194).

**Figure 25:**
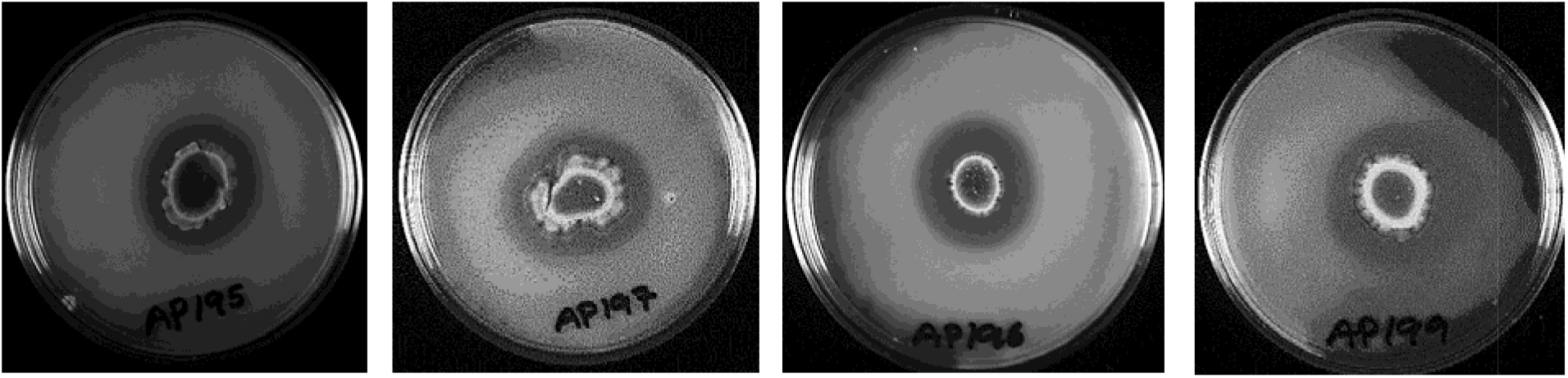
Pectate lyase activity of Bap strains (AP195, AP197, AP198, and AP199).

**Figure 26:**
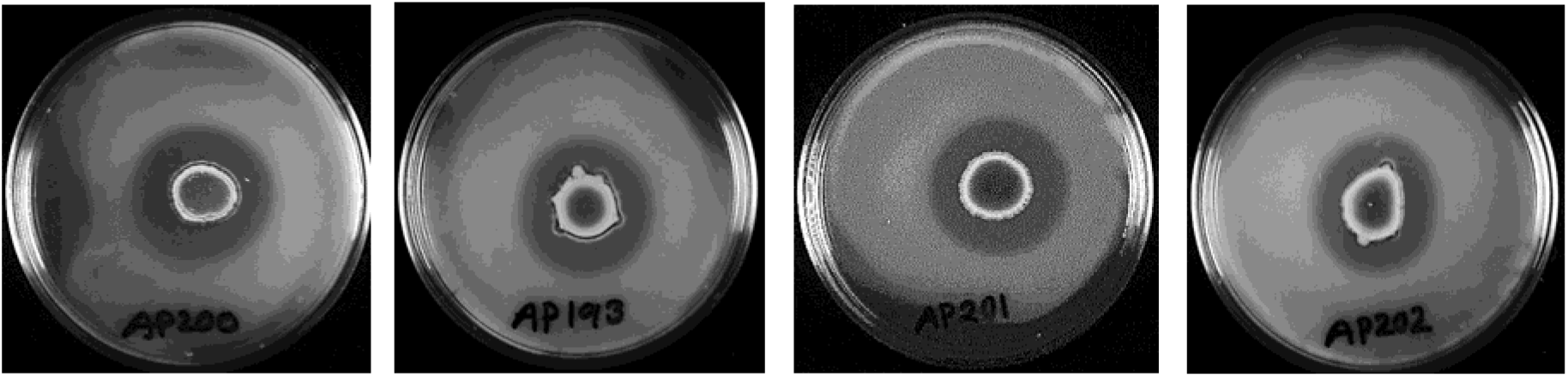
Pectate lyase activity of Bap strains (AP200, AP193, AP201, and AP202).

**Figure 27:**
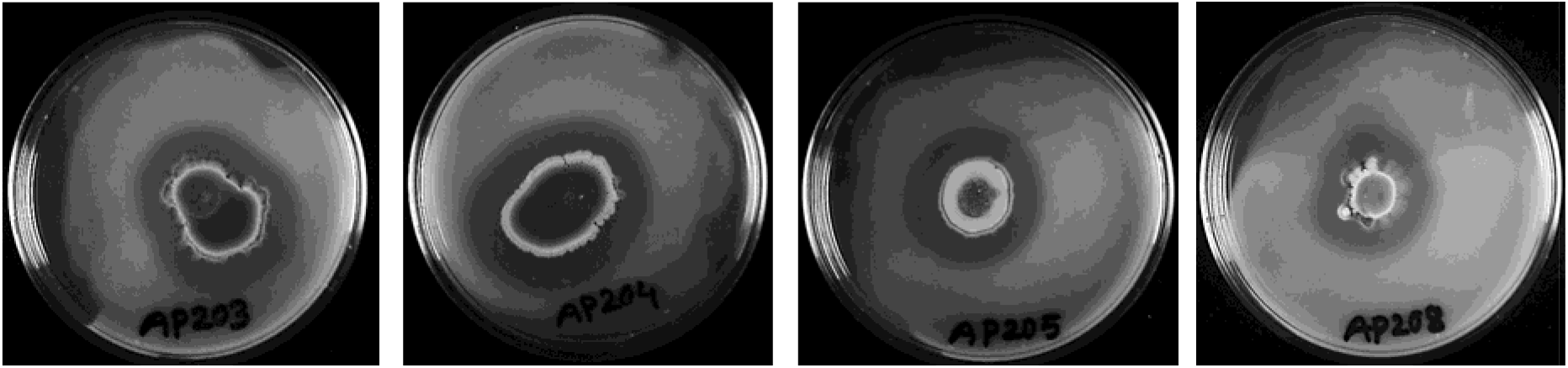
Pectate lyase activity of Bap strains (AP203, AP204, AP205, and AP208).

**Figure 28:**
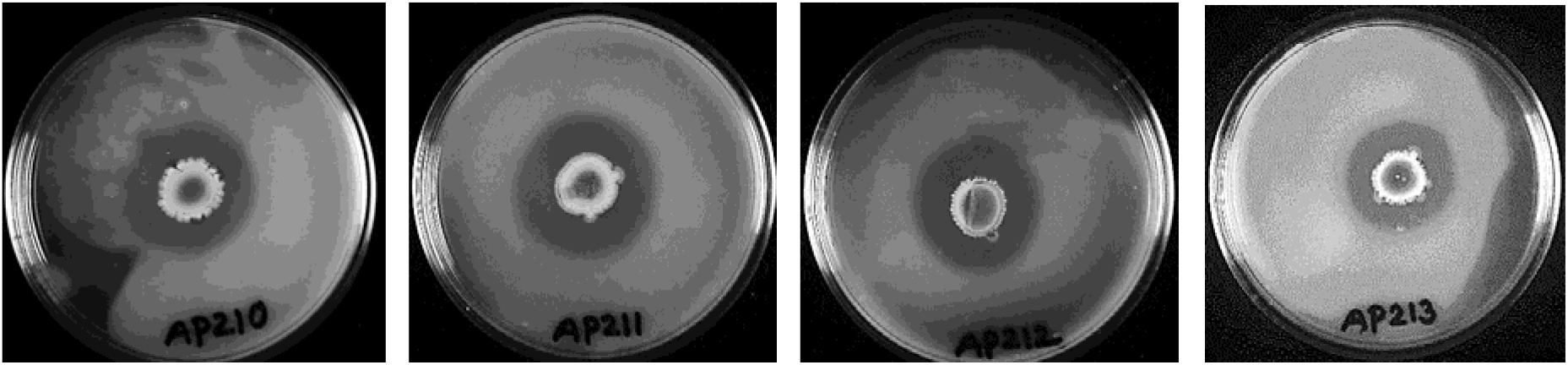
Pectate lyase activity of Bap strains (AP210, AP211, AP212, and AP213).

**Figure 29:**
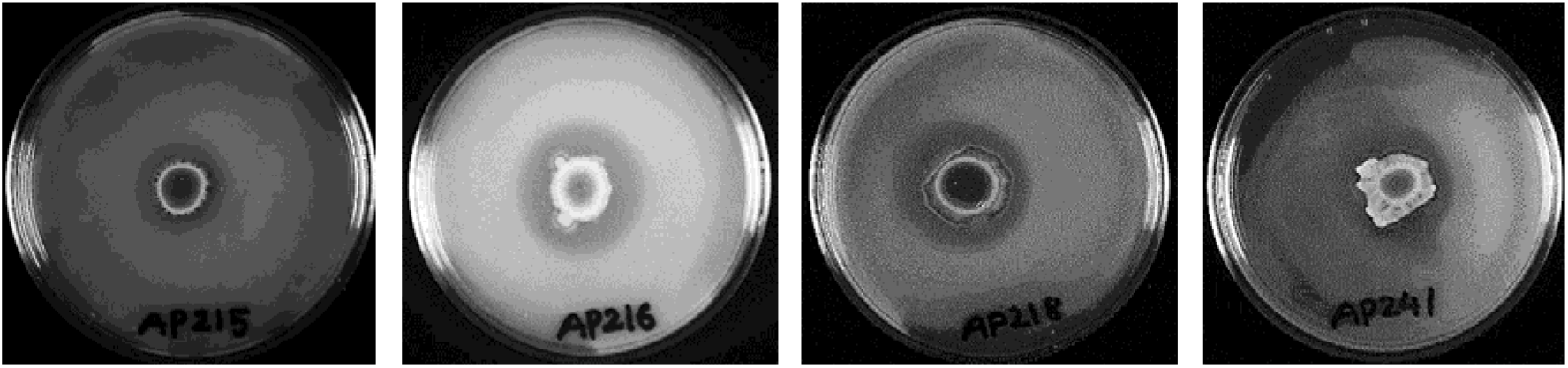
Pectate lyase activity of Bap strains (AP215, AP216, AP218, and AP241).

**Figure 30:**
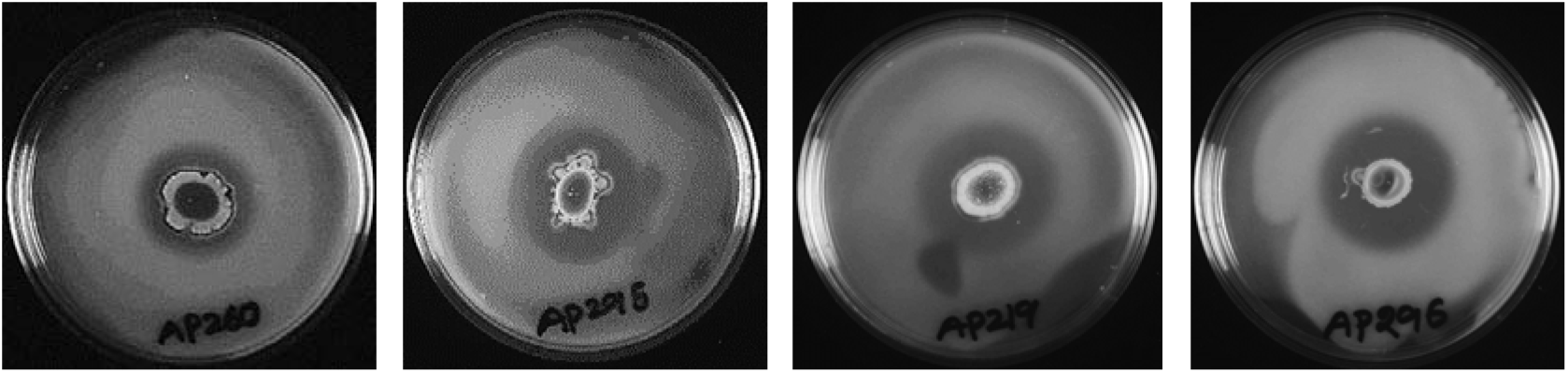
Pectate lyase activity of Bap strains (AP260, AP295, AP219, and AP296).

**Figure 31:**
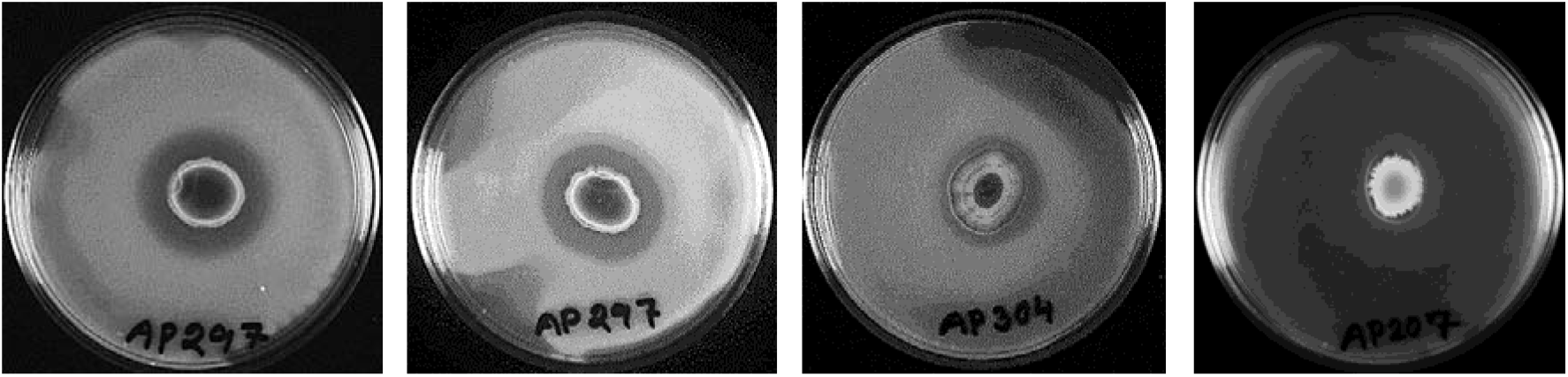
Pectate lyase activity of Bap strains (AP298, AP297, AP304, and AP297).

**Figure 32:**
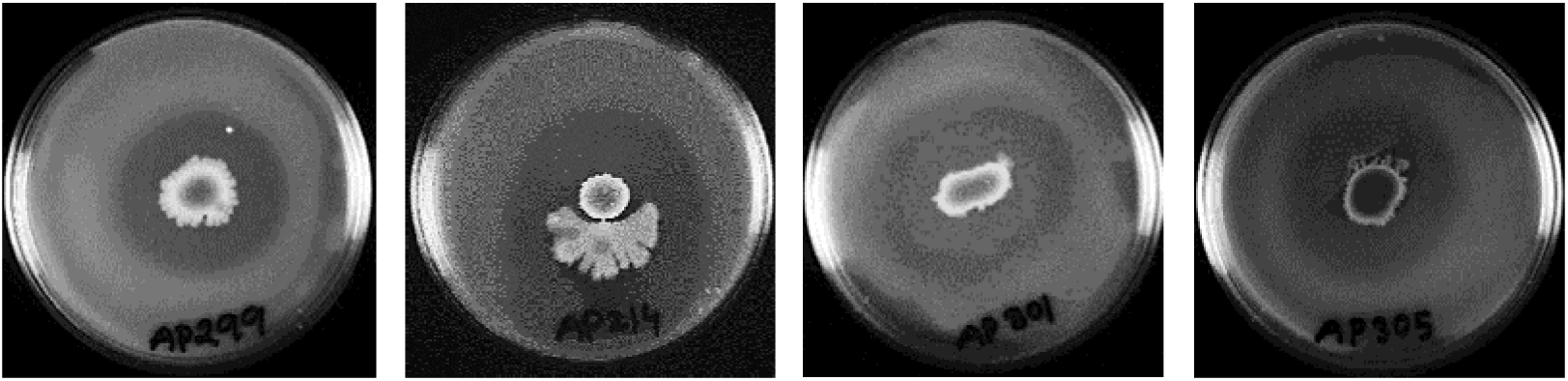
Pectate lyase activity of Bap strains (AP299, AP214, AP301, and AP305).

**Figure 33:**
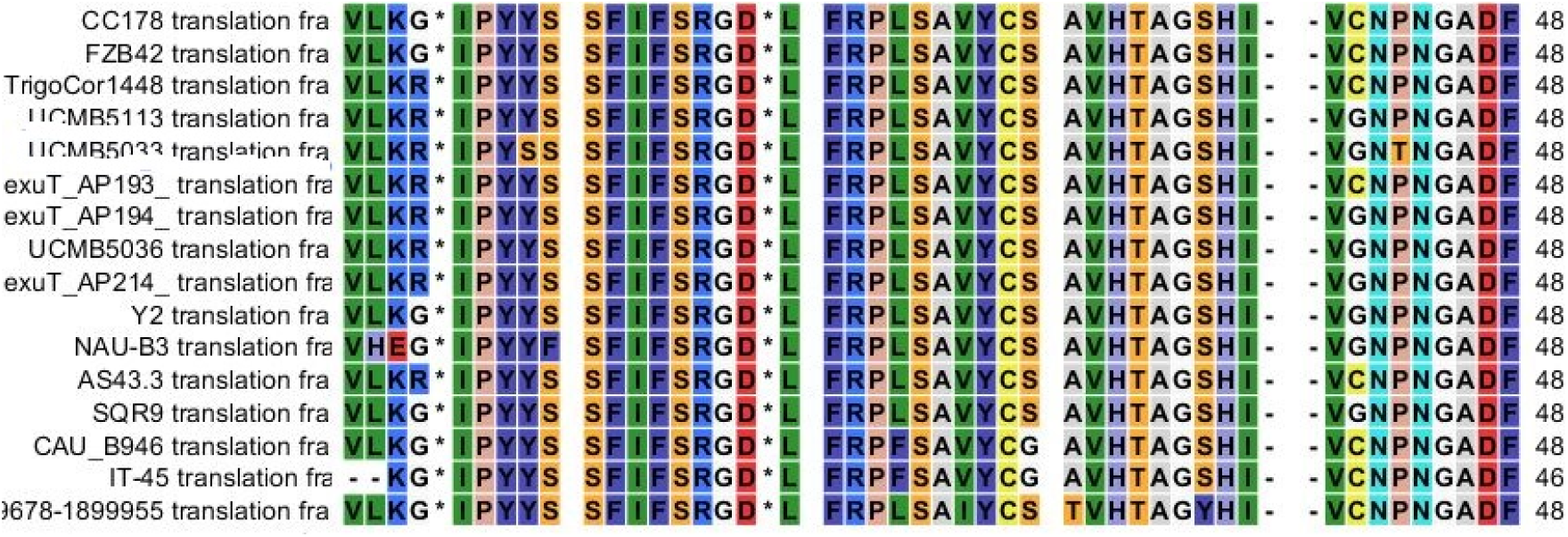
Multiple amino acid alignment of *uxuB* gene (AP193, AP194, and AP214) with reference strains.

**Figure 34:**
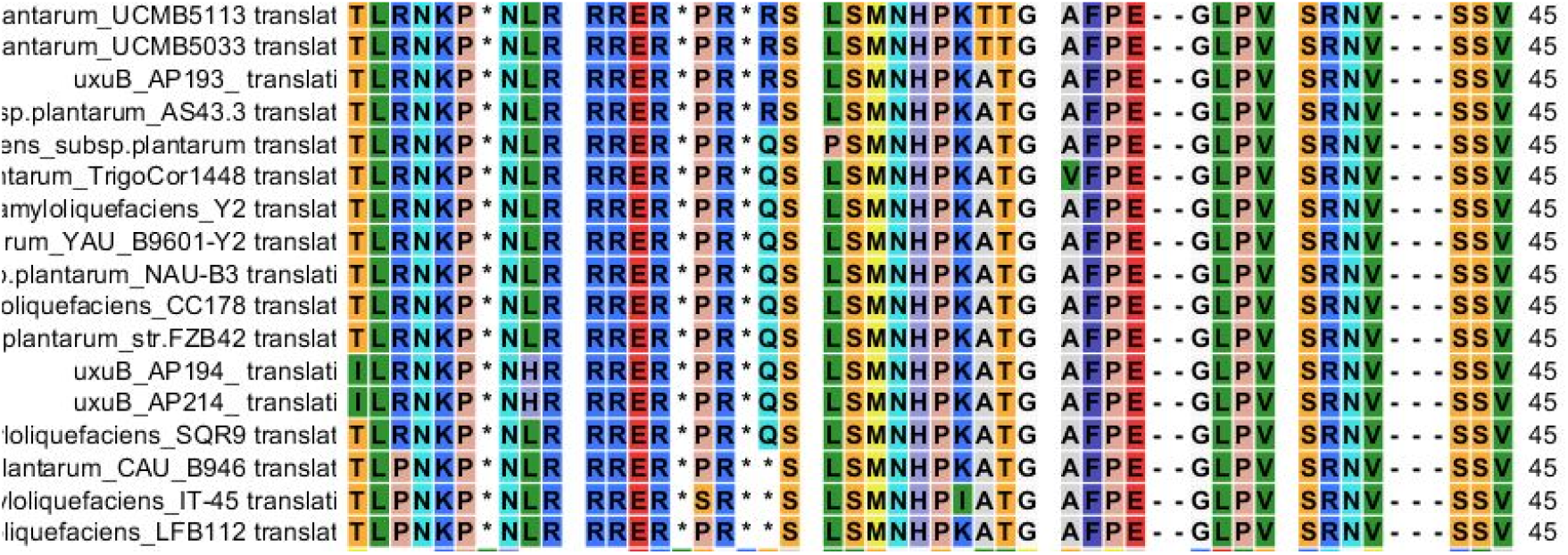
Multiple amino acid alignment of *exuT* gene (AP193, AP194, and AP214) with reference strains.

